# Disrupted RGS coupling underlies a clinically distinct subset of GNAO1-related disorder

**DOI:** 10.64898/2026.10.27.762269

**Authors:** Elizabeth G. Obarow, Erika J. Axeen, Cornelius F. Boerkoel, Joseph Gordon, Larissa Heideman, Hyun Lee, Laura van de Pol, Ashley Rice, Yibo Si, H. Dalton Taylor, Adam Tengölics, Chloe E. Viviers, Paul Slesinger, Henrik G. Dohlman

## Abstract

Heterotrimeric G proteins are transducers for a wide variety of neurotransmitter receptors. Mutations in the G protein α_o_ subunit are implicated in an early-onset neurological condition, *GNAO1*-related disorder, that manifests as movement disorders, developmental delays, and/or epileptic seizures. In a screen of disease-linked mutations in Gα_o_, we identified a subset that had no obvious biochemical defects, warranting further investigation. By structural modeling, protein-protein binding experiments in vitro and in cells, as well as measures of downstream ion channel electrophysiology, we identified a subset that impairs interactions with regulator of G protein signaling (RGS) proteins. RGS proteins accelerate G protein GTPase activity and are therefore critical for proper neurotransmitter signaling. Further, synthesis of current clinical data revealed that patients harboring these variants have comparatively mild symptoms. Our study demonstrates the importance of RGS regulation in G protein function, cell signaling, and human neurobiology.

## Introduction

G protein signaling is widespread in biology and mediates diverse processes, ranging from nutrient sensing in yeast to neuron firing in humans (*1*, *2*). When stimuli such as light, odors, hormones, or neurotransmitters reach the cell surface, G protein coupled receptors (GPCRs) detect these inputs, undergo conformational changes, and recruit heterotrimeric G proteins. G proteins, which consist of a Gα subunit and a Gβγ obligate dimer, are molecular “switches” that cycle between “on” and “off” states. In the “off” state, the G protein α subunit is bound to GDP. Upon agonist binding to a GPCR, the Gα subunit exchanges GDP for GTP and triggers the release of Gβγ. This characterizes the “on” state, where GTP-Gα and free Gβγ subunits go on to independently activate downstream effectors [reviewed in (*3*)]. To terminate signaling, GTPase-accelerating proteins (GAPs), such as regulator of G protein signaling (RGS) proteins, interact with Gα to accelerate hydrolysis of GTP, returning the G protein to the inactive, GDP-bound, heterotrimeric state [reviewed in (*4*)].

Alterations in G protein signaling have been implicated in cancer, addiction, and neurological disorders (*5–7*). In 2013, *de novo* heterozygous variants in *GNAO1* (encodes the Gα_o_ subtype) were first identified in four patients with epileptic encephalopathy (*8*). Since then, more than 200 unique variants of *GNAO1* have been reported in the ClinVar database. Here, we use the term *GNAO1*-related disorder (*GNAO1*-RD), for which there are three phenotypes: (i) infantile-onset developmental and epileptic encephalopathy (DEE) ± movement disorder, (ii) infantile- or early childhood movement disorder and neurodevelopmental disorder ± (non-DEE) epilepsy, and (iii) later childhood- or adult-onset movement disorder with variable developmental delay and intellectual disability (*9*).

To establish the mechanistic consequences of the Gα_o_ mutations, we and others have been investigating the ability of each mutant to interact with receptors and Gβγ, to stimulate an ion channel effector, and to bind and hydrolyze GTP (*10–14*). Our previous efforts identified four mechanistically distinct groups with differential signaling impacts: “Group 1” mutants exhibit sustained receptor association; “Group 2” mutants exhibit sustained heterotrimer assembly even when bound to GTP, and “Group 4” mutants express poorly in cells (*10*). In contrast, “Group 3” mutants exhibit no obvious biochemical defect; they have normal catalytic activity and are fully functional with respect to GPCR and Gβγ interactions. Structural modeling indicates that these “Group 3” mutations are located on the surface of the Gα_o_ subunit. Thus, it seemed likely that they disrupt interactions with a binding partner, but one other than GPCRs and Gβγ subunits. Here we consider their ability to bind RGS proteins.

Previous work has demonstrated the importance of RGS proteins in human physiology. Mutations in RGS9, or its anchoring protein R9AP, were identified in patients with bradyopsia, which is a rare condition that affects vision by impairing adaptation to changes in light (*15*). RGS9 is the predominant RGS subtype in the retina (*16*, *17*), and is required for the deactivation of phototransduction. That landmark study was the first to associate reduced RGS activity to a human disease. Gα_o_ is the predominant G protein subtype in the brain and is regulated by multiple RGS proteins that are expressed in a region-specific manner (*18*). Through our comprehensive approach, we identified a distinct set of mutations in Gα_o_ that disrupt interactions with multiple RGS protein subtypes. These new findings demonstrate how reduced RGS sensitivity may lead to a neurological disorder.

## Results

### *Gα_o_ mutants sustain proper receptor and G*βγ *coupling in cells*

To determine the mechanistic basis of *GNAO1*-RD, we previously performed a comprehensive functional screen of 55 Gα_o_ variants identified in patients (*10*). While most of the mutants exhibited a clear defect, a subset appeared to be fully functional with respect to enzyme activity, receptor coupling and subunit dissociation. Subsequent to the original screen, an additional five candidate Group 3 mutations were identified in ClinVar (S207F, E217D), in the literature (G184S, E208Q) (*19*), and in a patient seen by one of us (I185F) (C.F.B.). Here, we set out to establish the function of the newly identified Group 3 Gα_o_ mutants.

First, we measured the ability of mutants to interact with GPCRs and Gβγ in mammalian cells. To that end, we used the TRUPATH bioluminescence resonance energy transfer (BRET) method (*20*). In these experiments, the Gα subunit is tagged with a bioluminescent donor, *Renilla* luciferase (RLuc), and the Gγ subunit is tagged with a fluorescent acceptor, GFP. In the inactive heterotrimeric state, the BRET signal is high due to the close proximity of RLuc and GFP. In the active state, Gα and Gβγ are dissociated from each other and the BRET signal is diminished (**Fig. 1A**). Thus, we can measure the extent of G protein activation, over a range of agonist concentrations. We verified that the Gα_o_ mutants express and form heterotrimers by quantifying RLuc and GFP/RLuc (basal BRET) counts, respectively, before agonist addition (**Fig. S1A-B**). We also confirmed expression of the untagged mutant proteins by western blotting (**Fig. S1C-D**). The five new mutants showed robust activation in response to agonist, with some variation in the maximum effect (E_max_) relative to wild-type (**Fig. 1B, Data File S1**). As a control, we performed the same experiment with the previously identified Group 3 mutants (*10*), with similar results. Based on this analysis, we concluded that all eleven mutants interact properly with GPCRs and Gβγ.

**Figure 1.**
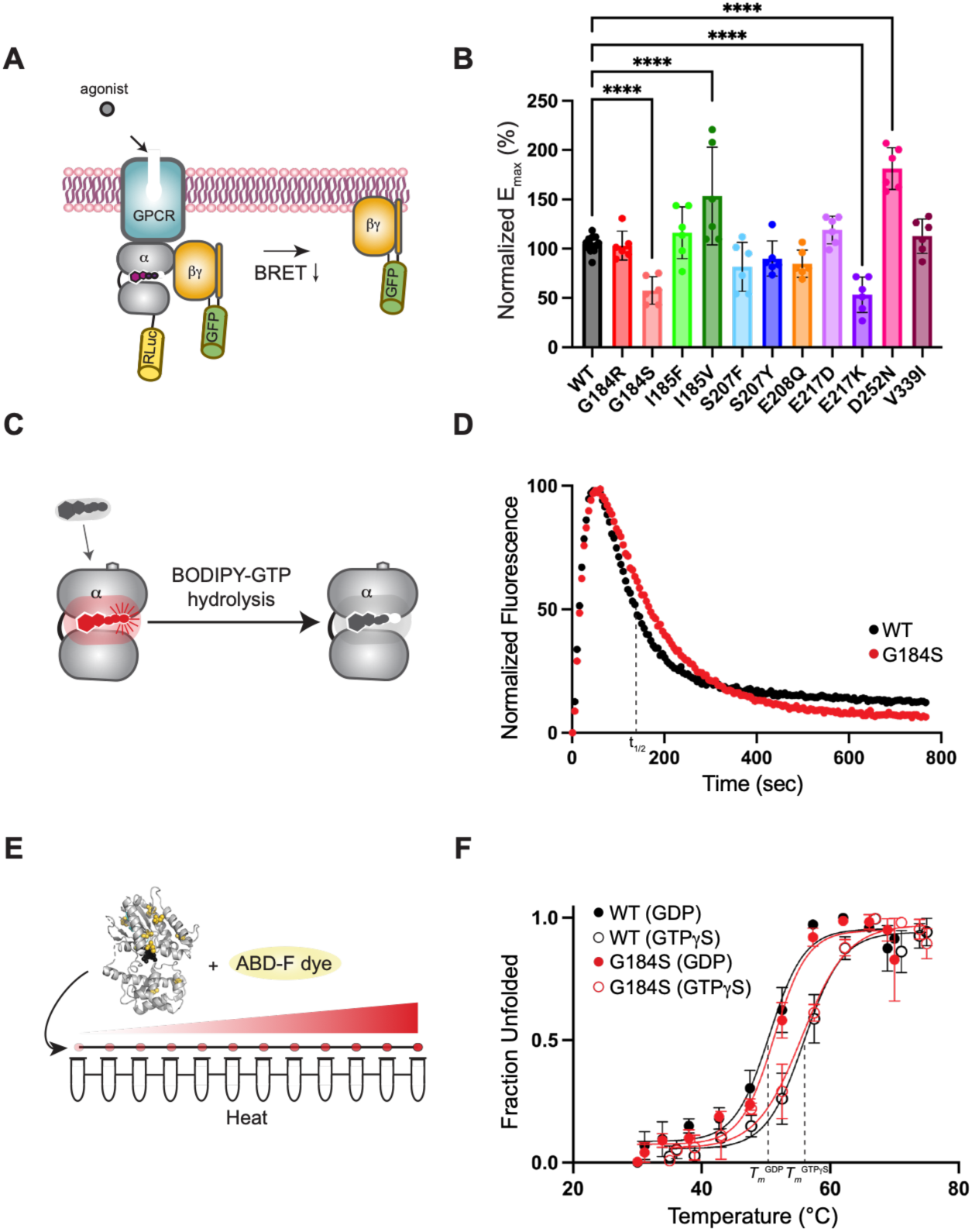
Activation, stability, and enzymatic activity of Gα_o_ disease mutants. (A) Agonist-induced G protein activation in the TRUPATH BRET system. (B) Normalized maximum effect (E_max_) for the Gα_o_ disease mutants upon activation of the μ-opioid receptor with DAMGO in HEK293T/17 cells. Data are means ± SD from three independent experiments, two measurements each. Statistical significance was determined by one-way ANOVA with Dunnett’s multiple comparisons test (α=0.01, ****p<0.0001). (C) GTP binding and hydrolysis of purified wild-type Gα_o_ bound to fluorescent GTP (BODIPY-GTP). (D) Representative GTP hydrolysis trace for wild-type (black) and mutant (G184S, red). Data are reported as means and are representative of two independent experiments, three measurements each. (E) Thermostability of purified Gα_o_ protein by fast determination of quantitative cysteine reactivity (fQCR). (F) Representative fQCR measurements for wild-type (black) and mutant (G184S, red). GDP-bound; closed circles. GTPγS-bound; open circles. Data are reported as means ± SD and are representative of two independent experiments, three measurements each.

### Gα_o_ mutants are functional as enzymes

To further establish the functionality of the Gα_o_ mutants, we performed fluorescence-based GTP hydrolysis and protein thermostability experiments. To measure enzymatic activity, we combined purified Gα protein with fluorescently labeled GTP (BODIPY-GTP) (*21*). Fluorescence increases upon BODIPY-GTP binding to Gα and subsequently decreases as Gα hydrolyzes the substrate to BODIPY-GDP (**Fig. 1C**). As with the previously identified Group 3 mutants, all five of the new Gα_o_ mutants bound and hydrolyzed GTP in a manner similar to wild-type (**Fig. 1D, table S1, Data File S2**). These findings indicate that the disease variants are stable and functional as enzymes.

However, we previously identified mutations that do not impact intrinsic enzymatic activity yet prevent Gα_o_ from assuming the GTP-bound, or activated, conformation (*10*, *11*, *22*). To test if the Gα_o_ mutants bind properly to guanine nucleotides and undergo the corresponding conformational changes, we measured protein thermostability in the presence of GDP or GTPγS (*23*). In this experiment, purified Gα is exposed to increasing temperatures, and upon thermal unfolding cysteine residues become exposed and labeled with a dye (**Fig. 1E**). The emitted fluorescence is plotted as a function of temperature, and the melting temperature (*T_m_*) is determined from the inflection point of the curves. For wild-type, the *T_m_* when bound to GDP was lower by ∼ 6 °C (51 °C) when compared to GTPγS (57 °C) (**Fig. 1F**, black). For the five new Gα_o_ mutants, all had *T_m_* values comparable to that of wild-type, for both GDP- and GTPγS-bound states (**Fig. 1F, table S2, Data File S3**). This indicates that the mutants are stable, bind to GDP and GTPγS, are catalytically active, and undergo nucleotide-specific conformational changes.

### Gα_o_ mutations slow effector channel deactivation

In the experiments described above, we demonstrated that the mutants exhibit normal enzymatic activity and are properly activated by receptors. We next sought to determine if they are properly inactivated. To that end, we measured coupling to G protein inwardly rectifying potassium (GIRK) channels. These channels are directly gated by Gβγ subunits following stimulation of GPCRs (*2*). RGS proteins oppose receptor action by enhancing the intrinsic GTPase activity of Gα, which promotes the formation of the inactive Gαβγ heterotrimer (**Fig. 2A**) (*24*, *25*). When RGS interactions are impaired, GIRK channels deactivate more slowly due to the slower sequestration of Gβγ (*26*). Thus, GIRK channel activity can provide a sensitive and dynamic readout of G protein activation and inactivation, and by extension the ability of RGS proteins to promote G protein turnover.

**Figure 2.**
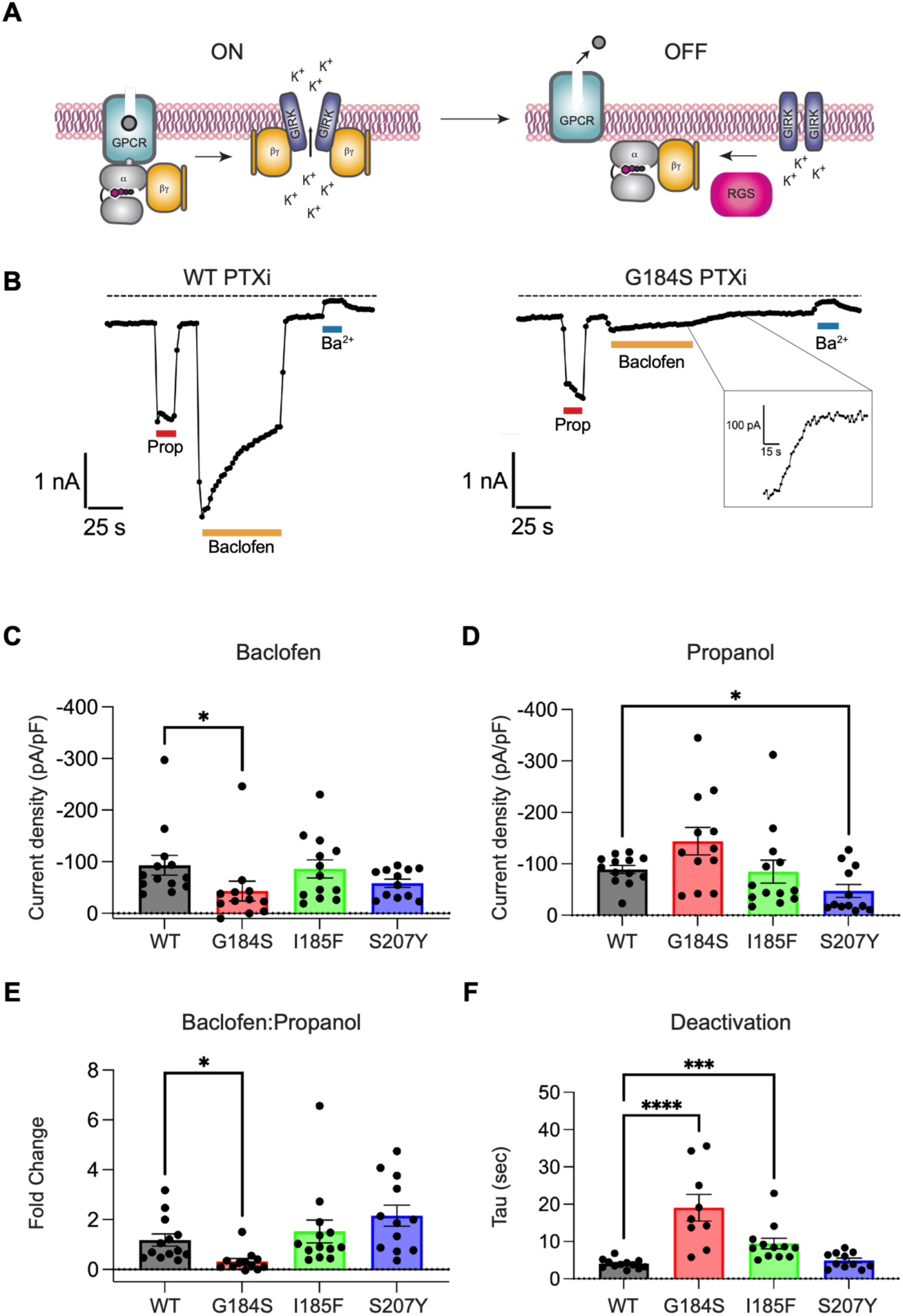
Downstream signaling outcomes of Gα_o_ disease mutants. (A) G protein mediated GIRK channel activation and inactivation. (B) Representative two-electrode voltage clamp (TEVC) recordings for HEK293T cells expressing GABA_B1/B2_, GIRK2, and either Gα_oPTXi_ WT-IRES-PTX_S1_ (left) or Gα_oPTXi_ G184S-IRES-PTX_S1_ (right) following treatment with propanol (Prop), baclofen, and barium (Ba^2+^). “PTXi”; pertussis toxin insensitive. (C) Baclofen current density for representative Gα_o_ mutants. Data are means ± SEM from 13 independent measurements for wild-type and I185F, and 12 independent measurements for G184S and S207Y. (D) Propanol current density for representative Gα_o_ mutants. Data are means ± SEM from 13 independent measurements for wild-type and I185F, and 12 independent measurements for G184S and S207Y. (E) Baclofen to Propanol ratio for representative Gα_o_ mutants. Data are means ± SEM from 13 measurements for wild-type and I185F, and 12 measurements for G184S and S207Y. (F) Tau deactivation for representative Gα_o_ mutants. Data are means ± SEM from 12 independent measurements for wild-type and I185F, 9 independent measurements for G184S, and 11 independent measurements for S207Y. Statistical significance in (C) and (D) was determined by one-way ANOVA (Kruskal-Wallis) with Dunn’s multiple comparisons test (α=0.05, *p<0.05). Statistical significance in (E) was determined by one-way ANOVA (Brown-Forsythe and Welch) with Dunnett’s T3 multiple comparisons test (α=0.05, *p<0.05). Statistical significance in (F) was determined by one-way ANOVA with Dunnett’s multiple comparisons test (α=0.05, ***p<0.001, ****p<0.0001). Only comparisons with statistically significant differences are notated.

To focus on ectopically expressed mutant Gα_o_ subunits, we used an established method to inactivate the endogenous Gα_i/o_ in HEK cells, which also express RGS proteins endogenously (*26*). Pertussis toxin uncouples Gα_i/o_ proteins from the receptor and prevents agonist activation of GIRK channels (*2*). A mutation in the C-terminal domain (C351A) renders Gα_o_ insensitive to pertussis toxin (PTXi). Under these conditions, the exogenous Gα_o_-PTXi, but not the endogenous Gα_o_, can sustain G protein activation of GIRK channels. We used this strategy to study the effects of three Gα_o_ mutations (G184S, I185F, S207Y), representing distinct residue positions at the RGS binding interface, in the background of the PTXi mutation.

We co-expressed the GABA_B_ receptor (R1/R2b) with GIRK2 and examined G protein activation via GABA_B_ receptor activation with baclofen, and G protein-independent activation of GIRK2 with propanol (*27*, *28*). We first verified in HEK cells that expressing the active protomer of pertussis toxin PTX_S1_ suppressed baclofen-activated currents, indicating uncoupling of the endogenous G protein with the GABA_B_ receptor (−2.94 ± 0.87 pA/pF, **Data File S4**). We then co-expressed wild-type Gα_o-PTXi_ to rescue GABA_B_ receptor activation of GIRK2 channels and observed reproducible baclofen-induced responses of ∼100 pA/pF (**Fig. 2B-C**). Propanol-activated currents for wild-type were also on average ∼100 pA/pF (**Fig. 2B, 2D**). We next examined the effects of expressing three representative Gα_o_ mutants (G184S, I185F, S207Y) on GIRK2 channel activation. While all three mutants supported some baclofen activation, the baclofen induced currents for G184S were significantly smaller (**Fig. 2C**). This was not likely due to reduced GIRK2 channel expression, since there was no significant difference in the propanol-activated current density for G184S (**Fig. 2D**) or the Ba^2+^-sensitive GIRK2 currents (**Fig. S2A**). Additionally, S207Y exhibited a reduction in propanol-induced but not in baclofen-induced currents (**Fig. 2C-D**). These results could indicate the mutant Gα_o_ is interacting with the channel and selectively interfering with alcohol gating. We have shown previously that Gα_i/o_ can interact directly with GIRK2 channels (*29*). Importantly, the baclofen-to-propanol ratio for G184S was significantly smaller than the ratio for wild-type (**Fig. 2E**), consistent with a change in G protein signaling (*30*). To address this, we examined the rate of channel closure (i.e., deactivation rate) after removal of baclofen. The presence of RGS is known to accelerate the deactivation rate by enhancing the GTPase activity of Gα_o_ to re-form the inactive Gαβγ heterotrimer (*4*). The rate of channel deactivation (τ) following removal of baclofen was significantly slower for G184S and I185F (**Fig. 2F**). Taken together, these results show impairments in G protein signaling with mutations in Gα_o_ that are predicted to alter RGS interactions, and that G184S, and to a lesser extent I185F, exhibit severe deficits.

### Gα_o_ mutations enhance agonist sensitivity

To corroborate our findings that some Gα_o_ mutants exhibited impaired regulation by RGS proteins, we examined if the functional effects of mutating these residues is conserved in other G proteins. The affected residues are widely conserved across the 16 Gα subtypes and perfectly conserved between human Gα_o_ and the yeast *Saccharomyces cerevisiae* Gα (Gpa1). Indeed, one of the new mutations described here (Gα_o_ ^G184S^) was instrumental in the discovery of the first RGS protein Sst2, in part because it phenocopied the loss of the *SST2* gene (*31*). Accordingly, we introduced the eleven mutations into Gpa1 and measured their effects on Gβγ-induced growth arrest and Gβγ-induced transcription in yeast (**Fig. 3A-B**) (*32*). For the growth arrest experiments, the mutants fell into two groups; one that responded like wild-type and the other that exhibited markedly larger zones of growth arrest (**Fig. 3C**, **S2A**). The latter phenotype indicates a greater extent of Gβγ release, possibly a consequence of impaired RGS action. For the transcription reporter assay we again saw two groups; whereas most of the mutants had wild-type-like responses, a smaller subset had left-shifted EC_50_s (**Fig. 3D, table S3, Data File S5**). A reduction in the EC_50_ indicates that G proteins are active at lower doses of agonist, potentially due to impaired RGS interactions (*32*). One outlier mutant (S207F; S325F in Gpa1) appeared fully functional in the growth assay but nonfunctional in the short-term transcription assay (**Fig. S3B)**. Taken together, the Group 3 mutations occur at positions conserved across the animal and fungal kingdoms, and across most of the 16 Gα subtypes; of these, a subset of mutants exhibit a defect resembling that of cells lacking a functional RGS protein.

**Figure 3.**
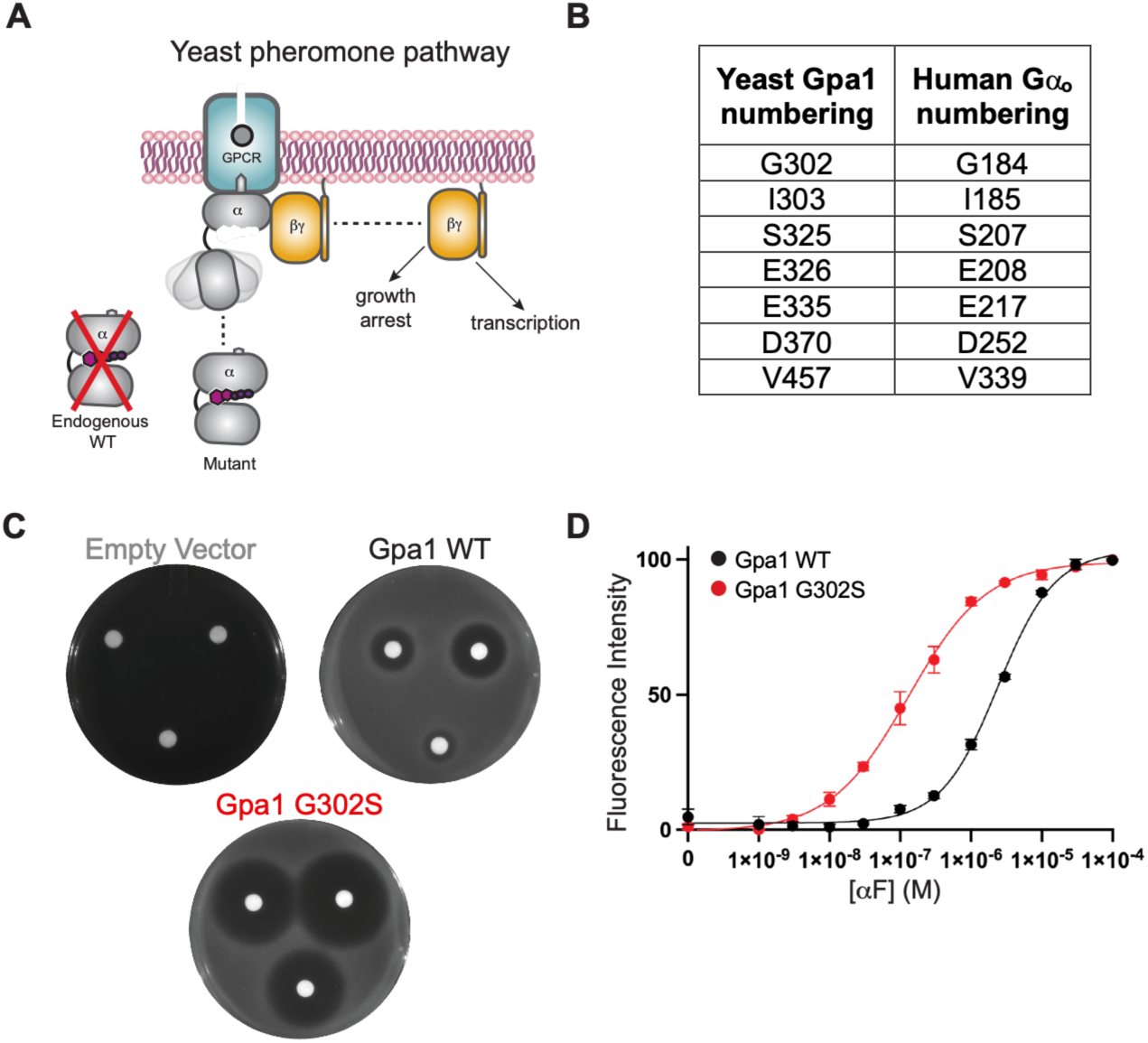
Evolutionary conservation of Gα_o_ disease mutants. (A) Gβγ-mediated growth arrest and transcription in the *Saccharomyces cerevisiae* pheromone pathway. Endogenous Gα wild-type is tagged with an auxin-inducible degron; when yeast are exposed to auxin the expression of wild-type is knocked down. (B) Comparison of yeast and human residue numbering system for the mutants of interest. (C) Representative growth inhibition outcomes for empty vector, wild-type, and Gpa1 G302S (corresponds to G184S in Gα_o_) after auxin treatment. Data are representative of two independent experiments, three measurements each. (D) Representative transcription response for Gpa1 wild-type (black) and mutant (G302S, red). Data are reported as means ± SD and are representative of two independent experiments, three measurements each.

### Gα_o_ mutants confer sustained subunit dissociation

Our GIRK and yeast experiments indicate that a subset of the disease mutations prolong the activated state of Gα_o_, likely by impairing its ability to be deactivated by RGS proteins. To test this, we performed a cell-based, kinetic BRET assay where a Gβγ effector fragment, G protein receptor kinase 3 c-terminus (GRK3ct), is tagged to the bioluminescent donor NanoLuc (NLuc), and Gβγ is tagged to the fluorescent acceptor Venus (*33*). When G proteins are activated, Gβγ is free and interacts with GRK3ct, emitting a high BRET signal (**Fig. 4A**). Conversely, antagonist addition causes Gβγ to reassociate with Gα, terminating Gβγ-GRK interactions and decreasing the BRET signal. When RGS is present, the deactivation rate is substantially faster (**Fig. 4B**) (*34*). Thus, the experiment measures time-dependent G protein activation and deactivation, enabling us to derive the rate constant of the deactivation phase (1/τ) for each of the mutants.

**Figure 4.**
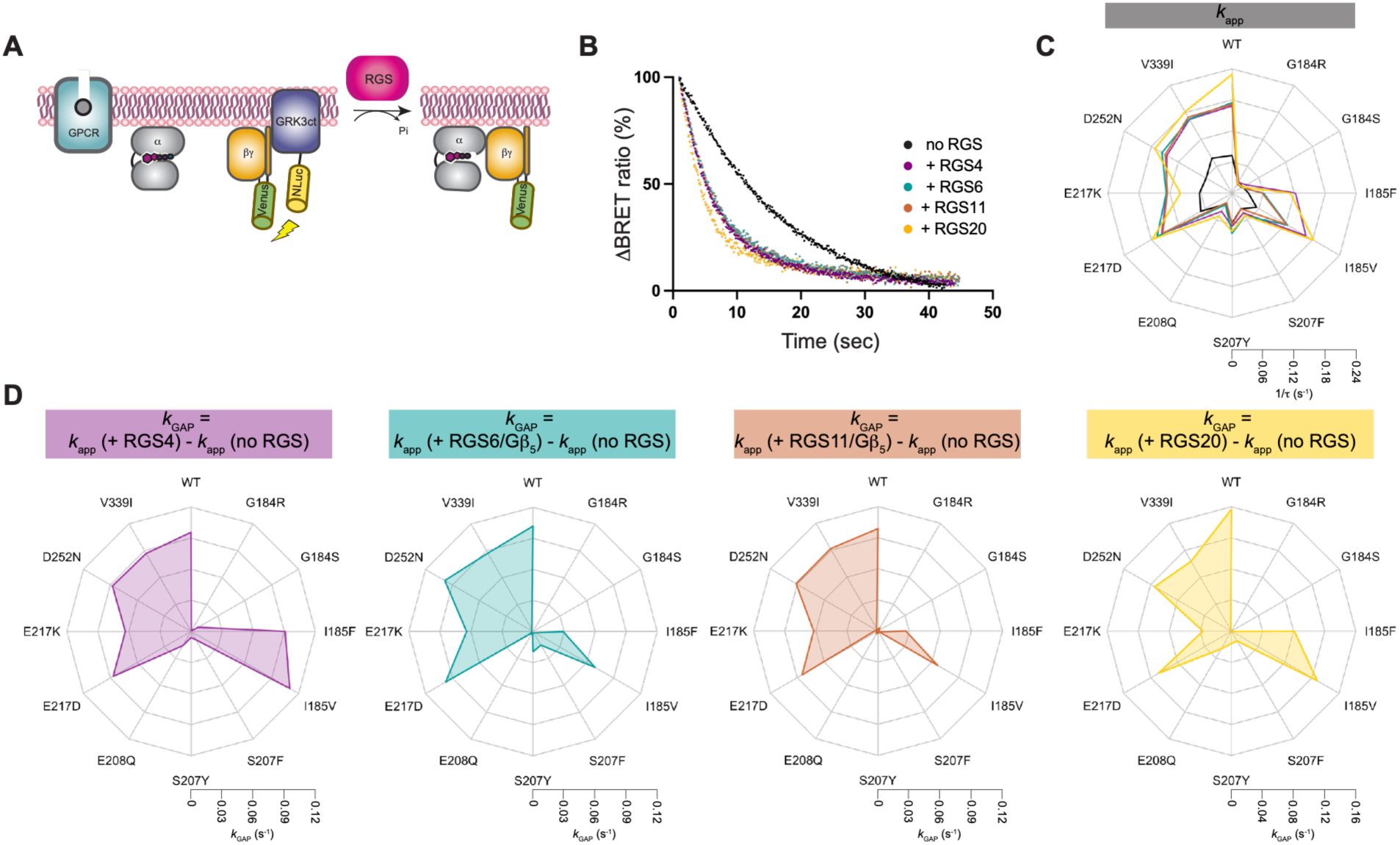
Deactivation of Gα_o_ disease mutants in human cells. (A) RGS-mediated G protein deactivation in the Venus-Gβ_1_γ_2_-masGRK3ct-Nluc BRET system. The μ-opioid receptor was activated with DAMGO and inactivated with naloxone in HEK293T/17 cells. (B) Representative deactivation phase trace for Gα_o_ wild-type without (black) and with (color) exogenous RGS proteins. Data are reported as means and are representative of three independent experiments, three measurements each. (C) 1/τ deactivation rate constants (*k*_app_) for Gα_o_ wild-type and mutants without an exogenous RGS protein. 1/τ values are means from three independent experiments, three measurements each. (D) Quantification of RGS activity for Gα_o_ wild-type and mutants with exogenous RGS proteins. *k*_GAP_ values were derived by subtracting the 1/τ deactivation rate constant without an RGS protein (*k*_app_) from the 1/τ rate constant with an exogenous RGS protein. *k*_GAP_ values are means from three independent experiments, three measurements each.

To establish a baseline of activity, we first measured the activation and deactivation kinetics of the mutants without ectopically expressed RGS proteins, quantified as *k*_app_ (**Data File S6)**. We derived *k*_app_ in the presence of two different Gα_o_-coupled receptors, the μ-opioid receptor (**Fig. S4B**) and the dopamine D2 receptor (**Fig. S4G**). Gα expression was verified by comparison with an empty vector control (**Fig. S4A**). As expected, the activation rates were unaffected by all but one of the mutants (**Fig. S4B**). Eight mutants (G184R/S, I185F/V, S207F/Y, E208Q, D252N) had deactivation rates that were substantially slower than wild-type (**Fig. 4C**, **Fig. S4B**), likely indicating impaired interactions with the RGS proteins that are endogenous in HEK cells. The remaining three mutants (E217D/K, V339I) had deactivation rates that were comparable to wild-type.

To determine the effect of specific RGS proteins on Gα_o_ mutant deactivation, we conducted the same experiment, this time in cells co-transfected with one of four RGS proteins (RGS4/6/11/20). These subtypes were selected because they show strong coupling to Gα_i/o_ (*34*) and are predominately expressed in the brain (proteinatlas.org) (*35*, *36*). To quantify the GTPase-accelerating protein (GAP) activity of RGS, we derived *k*_GAP_ values by subtracting the baseline rate constants (*k*_app_, no RGS) (**Fig. S4B**) from the rate constants with exogenous RGS (*k*_app_, +RGS) (**table S4, Fig. S4C-F, Data File S6**) (*37*). The presence of RGS4, RGS6, and RGS11 increased the deactivation rate for wild-type Gα_o_ by 2-fold (**Fig. 4C**, purple, blue, orange); RGS20 increased deactivation by 3-fold (**Fig. 4C**, yellow). As anticipated, most of the mutants with wild-type-like *k*_app_ values had wild-type-like *k*_GAP_ values, indicative of normal RGS-mediated deactivation (**Fig. 4D, table S5**). One mutant with a normal *k*_app_, E217K, appeared to be partially sensitive to RGS4, RGS6, and RGS11, but insensitive to RGS20. Conversely, most of the mutants with low *k*_app_ values in **Fig. 4C** had low *k*_GAP_ values relative to wild-type, indicative of impaired RGS-mediated deactivation. Three mutants exhibited intermediate or mixed sensitivities to exogenous RGS proteins. Based on the *k*_GAP_ values, I185F and I185V appeared to be partially sensitive to RGS4 and RGS20, and insensitive to RGS6 and RGS11. The remaining mutant, D252N, appeared to be sensitive to RGS4, RGS6, and RGS11, but partially sensitive to RGS20. Taken together, these experiments indicate that a majority of the Gα_o_ mutations impair the ability of one or multiple RGS proteins to terminate signaling in cells. The most severely impaired are G184R/S, S207F/Y, and E208Q; those showing partial impairment are I185F, I185V, and E217K (**Fig. 4D**).

### Gα_o_ mutants impair RGS binding

Our findings revealed a subset of Gα_o_ disease variants that appear to be refractory to RGS regulation. To establish the molecular basis for slowed G protein deactivation, we measured direct binding of RGS to Gα_o_ by surface plasmon resonance (SPR). In this experiment, a ligand (RGS) is immobilized onto a gold-plated sensor chip, and an analyte (Gα_o_) is flowed over the chip (**Fig. 5A**). The measured signal (in resonance units) is directly proportional to the magnitude of ligand-analyte interactions. We consider this method to be more reliable than co-precipitation experiments, which are not quantitative and cannot distinguish between poor binding to RGS and sustained binding to a competitor such as Gβγ. For these experiments, we used the RGS domain from RGS4. To ensure that the experiments were conducted in the high affinity state, we measured binding with and without aluminum fluoride (AlF_4_^-^). It has been previously established that RGS binds preferentially to Gα in the transition state for GTP hydrolysis (*38–41*). AlF_4_^-^ is a mimic for the pentavalent transition state that imposes a stable, semi-activated Gα for establishing RGS interactions in vitro.

**Figure 5.**
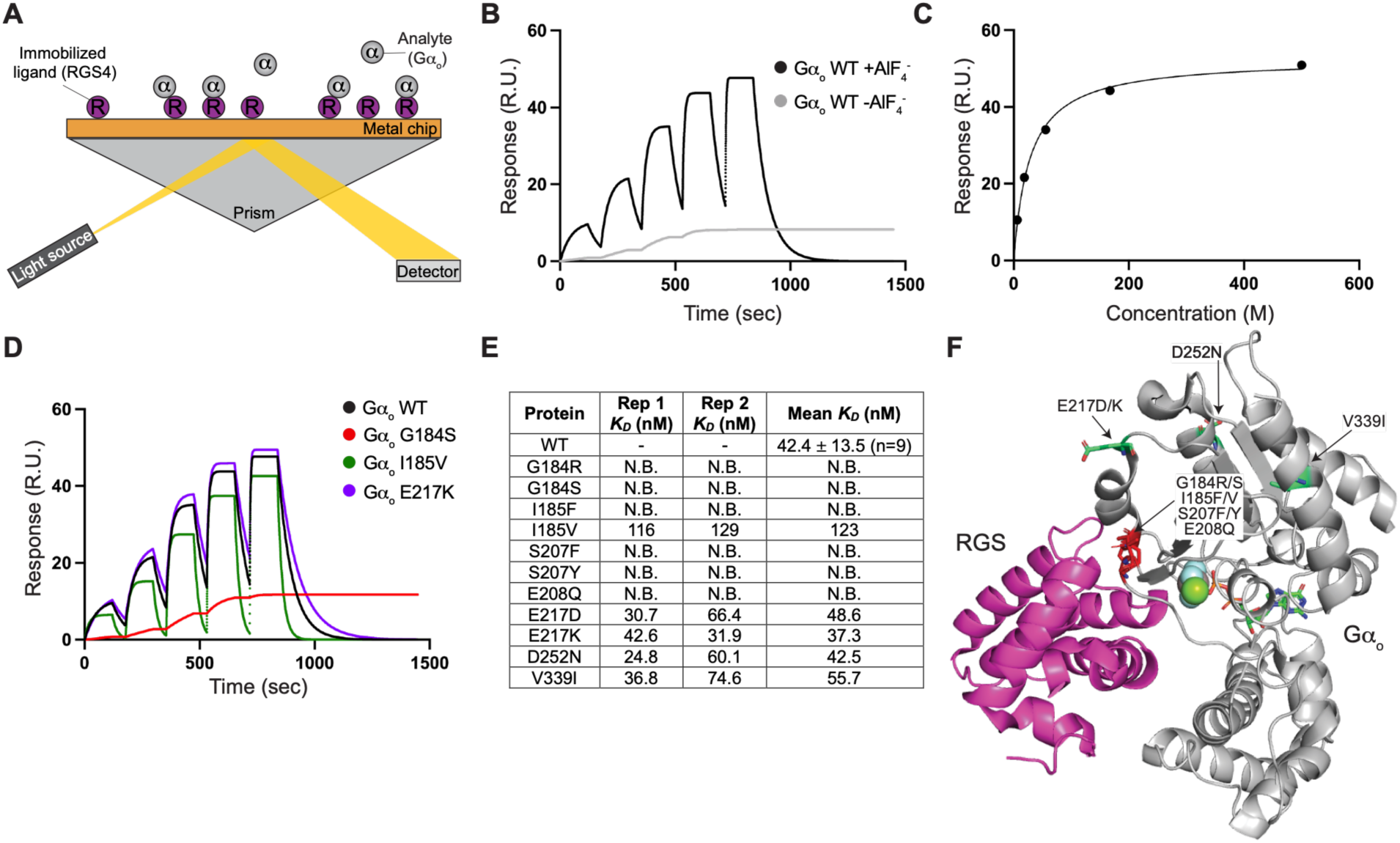
Gα_o_-RGS binding interactions in vitro. (A) Molecular interactions of Gα_o_ (analyte) and the RGS domain from RGS4 (ligand) by surface plasmon resonance. (B) Representative single-cycle kinetics of purified RGS interactions with purified Gα_o_ wild-type in the absence (gray) and presence (black) of aluminum fluoride (AlF_4_^-^). Data are representative of nine independent experiments. (C) Representative steady-state response curve for Gα_o_ wild-type. Data are representative of nine independent experiments. (D) Representative single-cycle kinetics of purified RGS interactions with purified Gα_o_ wild-type (black) and Gα_o_ mutants (color) in the presence of AlF_4_^-^. Data are representative of nine independent experiments for wild-type, two independent experiments for mutants. (E) Binding affinities of the purified Gα_o_ wild-type and mutants for RGS. “N.B.”, no binding. Data are from 9 independent experiments for Gα_o_ wild-type and 2 independent experiments for Gα_o_ mutants. (F) Structural modeling of the Gα_o_ residues that are mutated in *GNAO1*-related disorder. Red sticks; residues that impair RGS binding when mutated. Green sticks; residues that do not impair RGS binding when mutated. Gα_o_ subunit (light grey); RGS16 (purple); PDB: 3C7K.

We first performed a steady-state analysis to measure the binding of the RGS domain over a range of Gα_o_ concentrations. By this method, the affinity (*K*_D_) of wild-type Gα_o_-AlF_4_^-^ was 44.4 nM, which is consistent with the literature (**Fig. 5B-C**) (*42*). We also derived the kinetic affinities (k_a_/k_d_) and showed that they match the steady-state values (**table S6**). As expected, Gα_o_ failed to bind to the RGS domain in the absence of AlF_4_^-^. Of the mutants, six (G184R/S, I185F, S207F/Y, E208Q) had no detectable affinity, one bound poorly (I185V), and four (E217D/K, D252N, V339I) bound in a manner similar to that of wild-type (**Fig. 5D-E, Data File S7**). Correspondingly, the four residue positions that impaired RGS binding when mutated are located at the interface of Gα and RGS interactions (**Fig. 5F**), whereas the other three positions are not.

### Gα_o_ variants in human subjects

*GNAO1*-RDs are rare and present with a range of symptoms and severity. We obtained clinical information for patients with five of the seven variants that impaired RGS binding (**Table 1**; **Figure 5**). All the patients were classified as either infantile- or early childhood movement disorder and neurodevelopmental disorder ± (non-DEE) epilepsy (*9*). Two variants (S207F, S207Y) had been previously reported (*43*, *44*), two (G184S, E208Q) had been described in the literature (*19*) and were updated here, and one (I185F) had not been reported. The patient with the G184S variant has hypotonia and developmental delay without epilepsy or a movement disorder. The patient with the E208Q variant has diagnoses of intellectual disability, dystonia, and behavioral challenges.

**Table 1.**
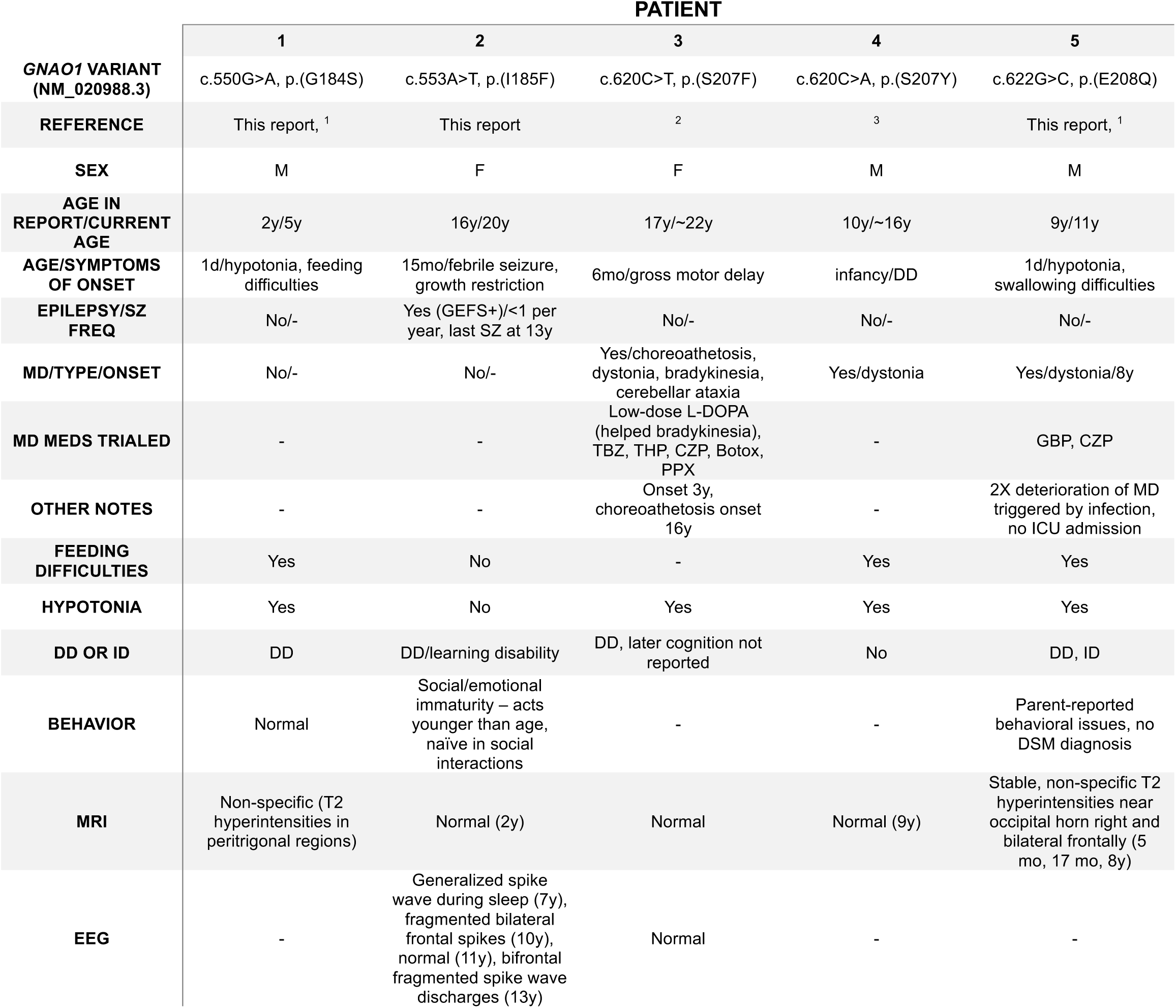
Clinical profiling of Gα_o_ disease mutants. GEFS+; Genetic Epilepsy with Febrile Seizures Plus. SZ; seizure. L-DOPA; levodopa. TBZ; tetrabenazine. THP; trihexyphenidyl. CZP; clonazepam. PPX; pramipexole. GBP; gabapentin. MD; movement disorder. DD; developmental delay. ID; intellectual delay. DSM; Diagnostic and Statistical Manual of Mental Disorders.

The patient with a novel variant (c.553A>T, p.I185F) is a 20-year-old female. She presented at 15 months of age with a febrile seizure and growth restriction. A head CT at 17 months and a head MRI at 2 years were unremarkable. She received speech therapy and was diagnosed at school age with learning disabilities in language, math, and memory. Notably, the patient exhibited social and emotional immaturities and poor impulse control. She continued to have generalized tonic-clonic seizures with and without fever. An EEG at 7 years of age showed generalized spike wave discharges, with subsequent EEGs at 10 and 13 years of age showing fragmented bilateral frontal spikes. Her syndrome was ultimately characterized as generalized epilepsy with febrile seizures plus (GEFS+). She was on valproic acid from ages 10-11 years but otherwise has remained off medication with no seizures since age 13. Whole exome sequencing sent in 2020 revealed the *GNAO1* variant. Thus, the patient is similarly mild in clinical manifestations of the others described in this study (**Table 1**) (*9*).

## Discussion

Since the discovery of *GNAO1*-RD in 2013, more than 200 missense mutations in Gα_o_ have been identified in patients. Previously, we characterized Gα_o_ mutants that exhibit sustained binding to receptors or Gβγ or both (*10*, *11*, *22*). Here, we investigated a set of Gα_o_ mutants that exhibit diminished binding to RGS proteins. Our mechanistic analysis included measures of G protein and GIRK channel deactivation, as well as direct measures of Gα-RGS binding. Further, we synthesized available clinical data of patients harboring the RGS-insensitive variants. Our analysis enabled us to identify a mechanistic link between RGS binding and human pathology.

Structural modeling revealed patterns between the location of the mutations and their effect on Gα_o_ function. All of the Group 3 mutations are located on the surface of the Gα_o_ subunit. The seven that cluster at the RGS binding interface (**Fig. 5F**) led to impaired deactivation for at least one of the tested RGS proteins in cells, and six substantially diminished binding to RGS in vitro. The remaining mutations (E217D/K, D252N, V339I) do not cluster at the RGS binding interface, had wild-type-like affinity for RGS, and were properly deactivated in cells. A notable exception was E217K, which bound normally in vitro but had reduced sensitivity to RGS in cells. Thus, it is still unclear how, or whether, these four mutations are associated with disease. A goal for the future is to test their ability to bind to candidate effectors (*45*), sustain proper cellular localization, or undergo post-translational modifications such as phosphorylation, palmitoylation, and myristoylation (*46*). Although the RGS-sensitive Gα_o_ mutations are not at known sites of modification, it is possible that they confer an indirect effect, such as altering a kinase or ubiquitin ligase binding site. On the other hand, it is possible that these variants are benign and unrelated to the patient phenotype.

RGS proteins were first discovered in yeast (*47*); cells lacking the *SST2* gene were shown to have increased pheromone sensitivity and defective recovery after signaling termination (*48*). Further desensitization screens revealed a mutation in the yeast Gα, Gpa1^G302S^, mimicked the loss of Sst2 (*31*); since the effects of the two mutations were not additive, it was inferred that Sst2 is a “regulator of G protein signaling”. Additional studies revealed that Sst2 acts specifically on Gpa1 (*49*, *50*) and accelerates its GTPase activity (*51*). Around this time, RGS proteins were also being identified and characterized in nematodes, plants and humans (*41*, *52–56*). Once the crystal structure of an RGS-Gα complex was solved in 1997 (*40*), the mechanism of action became clear. Specifically, RGS binds to Gα in the transition state for GTP hydrolysis, lowering the activation energy barrier by orienting catalytic residues in positions optimal for hydrolysis.

The discovery of the RGS-insensitive Gpa1^G302S^ mutation motivated the development of a heterozygous mouse model with the corresponding mutation in Gα_o_ (*Gnao1*^+/G184S^). One of the objectives of that effort was to determine the potential consequences of disrupting, either genetically or pharmacologically, RGS-Gα interactions (*57*). In these animals, the Gly-to-Ser substitution resulted in rare seizures, an increase in interictal epileptiform discharges, strain-dependent increased pentylenetetrazol kindling (as a measure of epileptogenesis susceptibility), and early death. Correspondingly, only one of the five patients described here has experienced any seizures, and they are rare. Thus, the *Gnao1*^+/G184S^ mouse provides striking evidence of a clear neurological deficit due to loss of RGS regulation of Gα_o_ and represents a bridge between our biochemical findings and patient outcomes.

The *GNAO1*-RD patients considered here represent a range of clinical manifestations (**Table 1**), with early-onset symptoms such as developmental delay and/or feeding issues, and varied manifestations of epilepsy and movement disorders (*9*). To date, they present with symptoms that are relatively less severe than those in other mechanistic groups (*10*), and lack epileptic encephalopathy or frequent severe dyskinetic crises. Accordingly, there are likely to be many patients with *GNAO1*-RD that are misdiagnosed or remain undiagnosed. As these new patients are identified, our mechanistic classification of disease variants may prove useful in identifying those most likely to benefit from treatments, including several currently in clinical trials (*13*, *58*, *59*).

Taken together, our work advances understanding of G protein signaling by demonstrating how disruption of negative regulation, rather than activation itself, can be sufficient to cause human neurological disease. Further, it provides direct mechanistic evidence linking defective RGS-G protein interactions to altered signaling dynamics in cells and functional consequences for neuronal regulation. Because RGS proteins are essential regulators of GPCR signaling throughout the nervous system, these findings have implications extending well beyond *GNAO1*-related disorders. They define a new class of signaling defects in which impaired signal termination, rather than defective activation, underlies disease.

## Materials and Methods

### Mutagenesis

Site-directed mutagenesis was performed using PrimeSTAR Max DNA Polymerase (Takara Bio; R045B). Primers were designed 5 bp 5’ and 20-30 bp 3’ to the relevant codon. See table S7 for primer sequences.

### TRUPATH bioluminescence resonance energy transfer (BRET) assays

G protein activation was measured in HEK293T/17 cells (ATCC; CRL-11268), as described previously (*10*, *20*). One million cells per well were seeded in 6-well plates (Corning; 3516) then transfected 6-8 hours later with plasmids encoding the μ-opioid receptor 1 (OPRM1), pcDNA5-Gα_o_-RLuc8, pcDNA3.1-Gβ_3_, pcDNA3.1-Gγ_9_-GFP2 in a 1:1:1:1 ratio and incubated at 37 °C. The next day (12-16 hours post-transfection), 80,000 cells/well were transferred to poly-D-lysine coated 96-well white clear bottom plates (Corning; 3903). The following day, cells were washed twice in assay buffer (1X Hank’s Balanced Salt Solution, 20 mM HEPES pH 7.4). Coelentrazine 400a (5 μM) (NanoLight Technology; #340) was added to each well and incubated in the dark for 5 minutes. Agonist (DAMGO, 30 μL of 30 μM; 10 μM final (highest concentration) and then seven 10-fold serial dilutions) was added to cells and incubated for another 5 minutes. Plates were read 4 times using spiral mode, and at room temperature using a CLARIOStar microplate reader (BMG LABTECH) (395 nm luminescence, 510 nm fluorescence). The BRET ratio was measured as the ratio of fluorescence over luminescence (GFP2/RLuc8). Net BRET was calculated by subtracting basal BRET (ratio at lowest dose of agonist) from the BRET ratio of each well.

### Immunoblotting

HEK293T/17 cells (ATCC; CRL-11268) were transfected with 200 ng pcDNA3.1-Gα_o_ wild-type and mutants and grown at 37 °C in Dulbecco’s Modified Eagle Medium (DMEM) (Corning; 10-013-CV) supplemented with 10% fetal bovine serum (Sigma-Aldrich; F2442) and 1% penicillin/streptomycin (Gibco; 15140122). After ∼40 hours, cells were lysed by sonication (Qsonica; Q500) on ice (5 sec on, 10 sec off, 30% amplitude, repeat twice) in 1X PBS pH 7, 150 mM NaCl, and 1% Triton-X-100, with Pierce Protease Inhibitor Tablets (Thermo Fisher; A32965), and centrifuged at 14,000 xg for 10 min. The supernatants were collected and resolved by 4-15% acrylamide gel electrophoresis and immunoblotting (20 mg each). Nitrocellulose membranes were blocked with 0.1% fish gelatin in 1X Tris-buffered saline with Tween-20 (TBST), probed with 1:500 Gα_o_ antibody A2 (Santa Cruz Biotechnologies, sc-13532, AB_2111645) and 1:5,000 mouse IgG Fc binding protein conjugated to Horseradish Peroxidase (m-IgG Fc BP HRP) (Santa Cruz Biotechnologies, sc-525409, AB_3101828), and then visualized with Clarity Western ECL Substrate (Bio-Rad, 1705061) on a ChemiDoc MP Imaging System (Bio-Rad). Quantification of Gα_o_ expression levels by fluorescence intensity was performed using ImageJ. For the loading control, blots were stripped with Restore Western Blot Stripping Buffer (Thermo Fisher; 21059) following manufacturer instructions. Blots were then re-probed with 1:5,000 vinculin polyclonal antibody (Proteintech; 26520-1-AP, AB_2868558) and 1:10,000 goat anti-rabbit conjugated to horseradish peroxidase (Jackson Immunoresearch; 111-035-045, AB_2337938) and visualized as explained above.

### Protein expression and purification

For production of Gα_o_, *E. coli* BL21(DE3) cells (NEBiolabs; C2527I) transformed with pET-SUMO-Gα_oA_ (wild-type or mutant) plasmid were grown at 37 °C in 2X Luria Broth to OD_600_ = 0.6. Protein expression was induced by addition of 600 μL of 1 M (600 μM final) isopropyl-β-D-1-thigalactoside (IPTG) and shaken at 18 °C overnight. For production of the RGS domain of RGS4 (Val51-Val178, 2243-2626), BL21(DE3) cells transformed with pET-SUMO-streptavidin binding protein-RGS4 (SBP-RGS) (*60*) plasmid were grown at 37 °C in 2X Luria Broth to OD_600_ = 0.6, then were moved to 25 °C for 1 hour before addition of 1 mL of 1 M (1 mM final) IPTG, and then shaken at 18 °C overnight. For protein purification, all steps were conducted at 4 °C. Cultures were centrifuged at 5400 xg for 30 minutes to collect cells and resuspended in G protein lysis buffer (25 mM H_2_KO_4_P, 25 mM HK_2_O_4_P pH 7, 300 mM NaCl, 500 μM TCEP, 50 μM GDP, 50 μM MgCl_2_) or RGS lysis buffer (50 mM HEPES pH 7.3, 300 mM NaCl, 5% glycerol, and 500 μM TCEP) + protease inhibitors (Thermo Fisher Scientific; A32965), lysed by sonication (Qsonica; Q500) on ice (8 sec on, 35 sec off, amplitude 100%, repeat 16X), and centrifuged at 36,635 xg for 30 minutes. DNA in the supernatant was precipitated with 300 μM of 5% polyethylenimine, and lysates were centrifuged at 2100 xg for 5 min. A column containing 3 mL (6 mL 50/50 slurry) HisPur Ni-NTA resin (Thermo Fisher Scientific; 88221) was prepared for protein binding by washing with 4 column volumes (CV) wash buffer I (25 mM H_2_KO_4_P, 25 mM HK_2_O_4_P pH 7, 300 mM NaCl, 500 mM TCEP, 50 μM GDP, 50 μM MgCl_2_) + 10 mM imidazole. Protein lysate was applied to columns. To wash columns, 4 CV of wash buffer I followed by 8 CV of wash buffer II (25 mM H_2_KO_4_P, 25 mM HK_2_O_4_P pH 7, 300 mM NaCl, 500 mM TCEP, 50 μM GDP, 50 μM MgCl_2_) + 30 mM imidazole were flowed through columns. To elute protein, 5 CV elution buffer (25 mM H_2_KO_4_P pH 7, 25 mM HK_2_O_4_P, 300 mM NaCl, 500 mM TCEP, 50 μM GDP, 50 μM MgCl_2_) + 250 mM imidazole were flowed through columns and collected. To cleave the SUMO tag, 0.1 mg of *Chaetomium thermophilum* SUMO protease (*61*) was added to eluates. To remove excess imidazole, protein was dialyzed overnight in purification buffer. The next day, the cleaved His6-SUMO tag was removed by flowing the dialyzed protein over the column packed with fresh Ni-NTA resin, and the flowthrough was collected. Protein was purified further using a HiTrap Q HP anion exchange chromatography column (Cytiva; 17115401) attached to an ӒKTA pure 25 FPLC (Cytiva). Briefly, 5 CV binding buffer (20 mM Tris pH 8) at 5 mL/min was run through the lines to prepare for protein binding. Protein (in binding buffer) was applied to the column at 1.5 mL/min. An additional 5 CV of binding buffer at 5 mL/min was run through the FPLC to wash out weak binding contaminants and ensure target protein binding. A gradient of 0-100% elution buffer (20 mM Tris pH 8, 1 M NaCl) in 45 min at 1.5 mL/min was performed and 1.5 mL fractions were collected. Pooled fractions containing Gα_o_ were concentrated and used for subsequent experiments.

### Nucleotide loading and hydrolysis

The ability of Gα_o_ to bind and hydrolyze GTP was determined using the fluorescent nucleotide, BODIPY FL GTP (Thermo Fisher Scientific, Invitrogen; G12411). Briefly, 50 nM of BODIPY FL GTP was equilibrated in GTPase buffer (20 mM Tris pH 8, 10 mM MgCl_2_) in a 1 mL cuvette. To initiate GTP binding and subsequent hydrolysis, 3 μM (final concentration) of purified Gα_o_ (bound to GDP) was added to the system. A Perkin Elmer LS50B spectrophotometer was used to measure fluorescence at room temperature (485 nm excitation, 530 nm emission).

### Thermostability experiments

Fast determination of quantitative cysteine reactivity (fQCR) experiments were conducted to measure protein thermostability, as described previously (*10*, *23*). To determine *T_m_* values, 10 μL of 1 mM GDP or GTPγS, 10 μL of 65 μM (3.25 μM final) purified Gα_o_ protein, 10 μL of 20 mM 4-fluoro-7-sulfamoylbenzofurazan (ABD-F) dye (Abcam; 91366-65-3), and 170 μL PBS pH 7 was added to PCR strip tubes. The samples were exposed to a 40-degree temperature gradient using a Biometra TAdvanced Thermal Cycler 96SG (Analytik-Jena) for 3 minutes. Fluorescence values were measured using a PHERAstar microplate reader (BMG LABTECH) (400 nm excitation, 500 nm emission).

### GIRK channel electrophysiology

G protein regulation of GIRK channels and the interaction with RGS was assessed using whole-cell patch-clamp electrophysiology with HEK293T cells, as described previously (*26*, *27*, *62*). For tissue culture, HEK293T (ATCC; CRL-1573) cells were maintained in Dulbecco’s Modified Eagle Medium (Life Technologies; 11965-118) supplemented with 100 U/mL penicillin, 100 mg/mL streptomycin, and 10% (v/v) heat-inactivated fetal bovine serum (HI-FBS; Thermo Fisher; 10438026). Lipofectamine 3000 (Thermo Fisher; L3000015) was used in accordance with the manufacturer’s protocol to transiently transfect cDNAs. To study the interaction of RGS with Gα_o_, we constructed a cDNA in pcDNA3 that expressed human Gα_o_ and the active protomer of pertussis toxin (PTX_S1_) to inactivate endogenous Gα_i/o_ proteins in HEK293T cells. The two genes are separated by an internal ribosomal entry site (IRES), which permits the expression of the two genes independently. In this way, we could ensure that each HEK293T cell that expressed Gα_o_ also expressed PTX_S1_. To make Gα_o_ resistant to PTX_S1_, we introduced a C351A mutation which does not affect Gα_o_’s function (*26*). We then engineered one of the following mutations, G184S, I185F or S207Y, into Gα_oPTXi_ to make wild-type Gα_oPTXi_-IRES-PTX_S1_, Gα_oPTXi_G184S-IRES-PTX_S1_, Gα_oPTXi_I185F-IRES-PTX_S1_, or Gα_oPTXi_S207Y-IRES-PTX_S1_ (*24*). All genes were cloned into pcDNA3 vector. For the negative control, HEK293T cells were transfected with cDNAs for mouse GIRK2 (100 ng) (*62*), GABA_B_R1b (100 ng), GABA_B_R2 (100 ng), eYFP (40 ng), and PTX_S1_ (100 ng). For testing mutants, HEK293T cells were transfected with cDNAs for mouse GIRK2 (100 ng) (*62*), GABA_B_R1b (100 ng), GABA_B_R2 (100 ng), eYFP (40 ng), and one of the Gα_oPTXi_-IRES-PTX_S1_ constructs (100 ng). 48-72 h post-transfection, cells were plated onto Poly-D-Lysine-coated (0.1 mg/mL) 12 mm glass coverslips (Electron Microscopy Sciences; 72290-03) in 24-well plates (VWR; 10062-896). Glass coverslips with cells were transferred to a 35 mm dish on a Nikon TE300 inverted microscope for electrophysiological recordings 1-48 h after plating. Whole-cell patch-clamp recordings were made as described previously (*26*, *27*, *62*). Transfected cells were identified by eYFP fluorescence. All data were acquired using an Axopatch 700B amplifier and pClamp 11 software (Molecular Devices). Whole-cell series resistance compensation was applied and used to estimate the cell size (pF). Patch-clamp experiments were conducted in 20K external solution containing 140 mM NaCl, 20 mM KCl, 2 mM MgCl_2_, 0.5 mM CaCl_2_ and 10 mM HEPES pH 7.4, (∼318 mOsm). Borosilicate glass electrodes (World Precision Instruments) had resistances of 3-4 MΩ and were filled with internal solution containing 20 mM NaCl, 130 mM KCl, 5.46 mM MgCl_2_, 5 mM EGTA, 2.56 mM K_2_ATP, 0.3 mM LiGTP and 10 mM HEPES pH 7.4 (∼300 mOsm). 100 mM PrOH, 100 µM baclofen and 1 mM BaCl_2_ were prepared in the external solution. Currents were recorded (10 kHz sampling and 1 kHz filter) from eYFP-positive cells using a voltage protocol of a holding potential of −40 mV, followed by a 60 ms step to −100 mV and then a continuous voltage ramp from −100 to 50 mV over 200 ms. Solutions were applied locally with a rapid perfusion pipet (Warner MP6 teflon manifold) and controlled by solenoid pinch valves (Warner Instruments), delivered in the following order: 20 K, 20 K + PrOH (100 mM), 20 K, 20 K + baclofen (100 µM), 20 K, 20K + BaCl_2_ (1 mM) and 20 K. The maximal inward current was measured at −100 mV. Average (mean) current was calculated during the 30 second drug application. The propanol-activated, baclofen-activated and Ba^2+^ inhibited current densities were calculated as (I_PrOH_ – I_Basal_)/pF, (I_Bac_ – I_Basal_)/pF, and (I_Basal_ – I_Ba2+_)/pF, respectively. To adjust for differences in GIRK2 expression, baclofen-induced activation was compared to G protein-independent propanol activation (*28*), and calculated as a ratio of baclofen-induced to PrOH-induced currents (I_Bac_ – I_Basal_)/(I_PrOH_ – I_Basal_). To measure the deactivation (τ), the decay of the current following washout of baclofen was fit with a single exponential in Clampfit. Cells not expressing a propanol-activated current were excluded from the analysis.

### Gβγ-induced growth arrest (halo) assay

A saturated culture of the *S. cerevisiae* BY4741 (MAT**a** *his3*Δ1 *leu2*Δ0 *met15*Δ0 *ura3*Δ0) strain with a V5 AID tag incorporated at *GPA1* (Kan) and *OSTIR1* incorporated at *LEU2* (pGPD1-osTIR1-tADH1::LEU2) (hereafter BY4741-GPA1-AID) was transformed with pRS316 empty vector, pRS316-GPA1 wild-type or pRS316-gpa1 mutants (Supplementary Materials). Cells were maintained at 30 °C unless otherwise indicated. Transformed yeast were grown to saturation for 40-48 hours in liquid selective medium. The next day, cells were mixed with 0.5% molten agarose in a 0.1:4 yeast:agarose ratio, spread on selective medium plates containing 10 μM (final concentration) indole-3-acetic acid (IAA) and exposed to 4 ng, 13 ng, or 40 ng synthetic α-factor on paper discs. Cells were kept dark, grown at room temperature, and imaged after 48 hours using a ChemiDoc MP Imaging System (Bio-Rad).

### Transcription reporter assay

A saturated culture of BY4741-GPA1-AID was transformed with pRS316 empty vector, pRS316-GPA1 wild-type or pRS316-gpa1 mutants and pRS423 FUS1-lacZ and grown in selective medium to saturation overnight. Cells were maintained at 30 °C unless otherwise indicated. The following day, the culture was back diluted twice (in the morning and afternoon), grown overnight, and either supplemented with 100 μM IAA (final concentration) + 0.3% DMSO or 0.3% DMSO only (control) and shaken until the cultures reached OD_600_ = 1.0 (∼60 minutes). Cells (90 μL) were added to a 96-well plate containing 10 μL of α-factor (0-100 μM final) and incubated for 90 minutes. Fluorescein di-β-D-galactopyranoside (FDG) solution (135 mM PIPES pH 7.2, 0.25% Triton, 0.5 mM FDG) was added to each well (20 μL/well) and the plate was incubated at 37 °C for 90 minutes. OD_600_ and fluorescence (485-15 excitation, 580-20 emission) was measured using a CLARIOstar Plus microplate reader (BMG LABTECH).

### Kinetic bioluminescence resonance energy transfer (BRET) assays

HEK293T/17 (ATCC; CRL-11268) cells were grown at 37 °C in DMEM with 1 mM sodium pyruvate (Corning; 10-013-CV), minimum Eagle’s medium non-essential amino acids (Gibco; 11140050), and 10% fetal bovine serum (Sigma-Aldrich; F2442). Cells (50,000/well) were transferred to a 96-well plate (Corning; 3903) in the above media supplemented with 0.1% Matrigel (Corning; 354234). Cells were transfected in a 1:1.5:1.5:0.5:2:X ratio of human μ-opioid receptor 1 (OPRM1), Venus 156-239-Gβ_1_, Venus 1-155-Gγ_2_, masGRK3ct-Nluc-HA, Gα_oA_, and different amounts of RGS construct (indicated in parentheses): RGS4 (50.4 ng), RGS6/Gβ_5s_ (4.2 ng), RGS11/Gβ_5s_ (25.2 ng), RGS20 (25.2 ng), with Lipofectamine LTX and PLUS (Thermo Fisher; 15338100). At 24 hours post-transfection, cells were washed and resuspended in BRET buffer (1X PBS pH 7, 0.5 mM MgCl_2_, 0.1% glucose). Furimazine (Nluc substrate) (Promega; N1110) was prepared according to manufacturer instructions and added to cells. BRET measurements were taken using the BRET1 plus module on a PHERAstar FS microplate reader (BMG LABTECH). All measurements were conducted at room temperature. To activate GPCR signaling, 50 μL of 200 μM (50 μM final) DAMGO or 50 μL of 400 μM (100 μM final) dopamine (agonists) was injected 10 seconds into the run. To deactivate GPCR signaling, 50 μL of 200 μM (50 μM final) naloxone or 50 μL of 400 μM (100 μM final) haloperidol (antagonists) was injected 45 seconds into the run. The ΔBRET ratio was determined by dividing the signal from Venus-Gβ_1_γ_2_ (535 nm ± 30 nm) by the signal from masGRK3ct-Nluc-HA (475 nm ± 30 nm) and normalized by subtracting the average baseline value from the experimental BRET signal values. Activation rate constants (1/τ) were derived by fitting a “plateau followed by one phase association” curve fit to the activation phase of the BRET curves (X0=10.9 sec). Deactivation rate constants (1/τ) were derived by fitting a one phase decay curve fit to the deactivation phase of the BRET curves. *k*_GAP_ values were derived by subtracting 1/τ (no RGS) by 1/τ (+ RGS). Radial plots in Figure 4D were made using R programming.

### Surface plasmon resonance

The day before the experiment, purified Gα_o_ was buffer exchanged and diluted to 10 μM into Gα SPR buffer (20 mM HEPES pH 7.3, 150 mM NaCl, 1 mM MgCl_2_, 1 mM TCEP, 0.05% Tween-20, 10 μM GDP, 30 μM AlCl_3_). Purified SBP-RGS was dialyzed into RGS SPR buffer (20 mM HEPES pH 7.3, 150 mM NaCl, 1 mM MgCl_2_, 1 mM TCEP, 0.05% Tween-20) and diluted to 10 μM. The day of the experiment, 1 mM NaF was added to a portion of the Gα SPR buffer. Wild-type Gα_o_ was split into two tubes and diluted to 500 nM in Gα SPR buffer ± NaF to form the AlF_4_^-^ complex and mimic the transition state for high affinity RGS binding. Gα_o_ mutants were diluted to 500 nM in Gα SPR buffer + NaF. SBP-RGS was diluted to 100 nM in RGS SPR buffer. To measure protein-protein binding, the single-cycle kinetics program on a Biacore 8K (Cytiva) was used. Briefly, 100 nM of SBP-RGS was immobilized on a Series S Sensor Chip SA (Cytiva) and Gα was flowed over the chip at 5 different concentrations in 3-fold increments (500 nM, 166.7 nM, 55.6 nM, 18.5 nM, 6.2 nM). Measurements were taken at ambient temperature. Kinetic values (*k_D_*, k_a_, k_d_) were obtained using the Biacore Insight Evaluation Software (Cytiva).

### Clinical analysis

This study includes five patients with *GNAO1* gene alterations and neurodevelopmental disorders. Data from two patients (S207F, S207Y) were obtained from the literature (*43*, *44*). Two patients (G184S, E208Q) were identified in a literature search (*19*), and the relevant author was contacted (by L.P.); these patients were evaluated by direct observation at Amsterdam University Medical Center. One patient with a novel variant (I185F) receives care by one of us (C.F.B.) and was evaluated by direct observation at the British Columbia Children’s Hospital. The family of the patient with the novel variant provided consent to be included in this study.

## Statistical analysis

Quantitation, generation of dose-response curves, curve fitting, and statistical analysis (SD, SEM, one-way ANOVA) was done using GraphPad Prism (GraphPad Software Inc., San Diego, CA). Biological replicates are independent, parallel measurements of biologically distinct samples obtained on different days. Technical replicates (duplicate or triplicate) are repeated measurements of the same sample and represent random noise associated with protocols or instrumentation. Ordinary one-way ANOVA was chosen for most statistical tests because more than two groups (wild-type vs. eleven Gα_o_ mutants) were being compared simultaneously. Dunnett’s multiple comparisons test was chosen for most post-hoc tests because the eleven Gα_o_ mutants were being compared against a single control (Gα_o_ wild-type). For the GIRK channel experiments, alternative one-way ANOVA and the corresponding post-hoc tests (Kruskal-Wallis with Dunn’s, Brown-Forsythe and Welch with Dunnett’s T3) were chosen because the assumptions of normal distribution and equal variances, respectively, were not made.

## Supplementary Materials

References (*60*)-(*62*)

Data Files S1-S7

## Acknowledgments

The authors wish to thank C. Travis, A. Tripathy, and B. Golitz (UNC-CH) for their technical and conceptual support. We thank Yan Cao (UF) for providing plasmids for the Gβγ-GRK BRET experiments.

## Author contributions

E.G.O., E.J.A., C.F.B., Y.S., P.S., H.G.D.: conceptualization. E.G.O., E.J.A., C.F.B., J.G., L.H., H.L., A.R., H.D.T., A.T., L.P., C.E.V., P.S., H.G.D.: formal analysis. P.S., H.G.D.: funding acquisition. E.G.O., J.G., A.R., H.D.T., A.T., C.E.V.: investigation. P.S., H.G.D.: project administration. P.S., H.G.D.: supervision. E.G.O., E.J.A., C.F.B., J.G., L.H., H.L., A.R., H.D.T., A.T., L.P., C.E.V., P.S., H.G.D.: validation. E.G.O., E.J.A., J.G., A.T., P.S., H.G.D.: writing - original draft. E.G.O., E.J.A., C.F.B., L.H., H.L., L.P., P.S., H.G.D.: writing - review & editing.

## Data and materials availability

TRUPATH BRET reagents are available on AddGene (Kit #1000000163). All other unique reagents generated in this study will be available upon request. Data reported in this paper will be shared by one or more of the corresponding authors upon request. The study did not generate new unique code. All other data needed to evaluate the conclusions in the paper are present in the paper or the Supplementary Materials.

---

**Figure S1.**
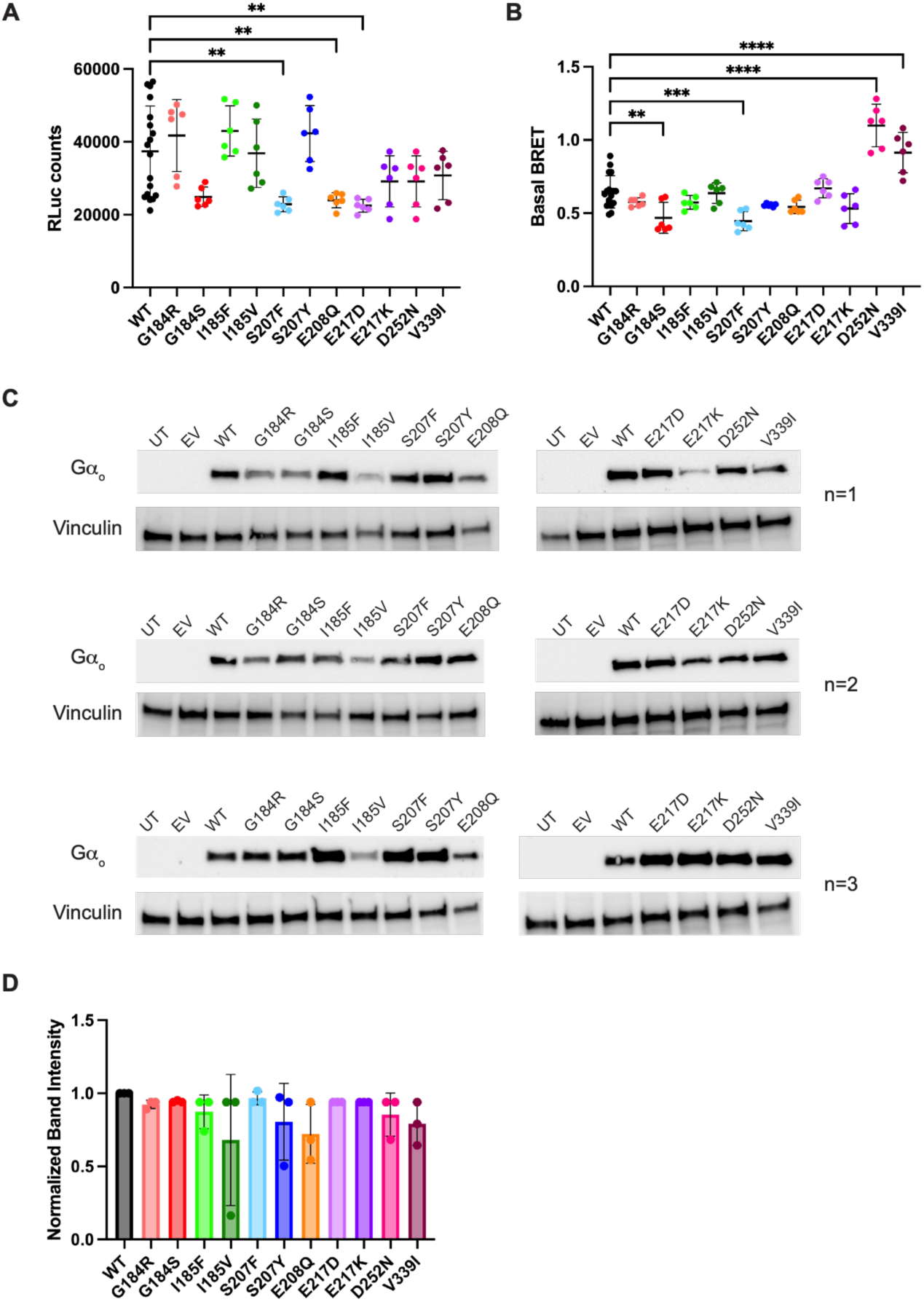
RLuc counts, basal BRET, and immunoblots for Gα_o_ wild-type and mutants. (A) RLuc counts for the Gα_o_ disease mutants tested by TRUPATH BRET, as a measure of Gα_o_ expression in HEK293T/17 cells. Data are means ± SD from three independent experiments, two measurements each. (B) Basal BRET ratio (GFP/RLuc) counts for the Gα_o_ disease mutants tested by TRUPATH BRET, as a measure of heterotrimer formation in HEK293T/17 cells. Data are means ± SD from three independent experiments, two measurements each. Statistical significance was determined by one-way ANOVA with Dunnett’s multiple comparisons test (α=0.01, **p<0.01, ***p<0.001, ****p<0.0001). (C) Immunoblots of untransfected (UT) and transfected HEK293T/17 cells with pcDNA3 Gα_o_ wild-type and mutants, blotted for Gα_o_ (A2), and corresponding vinculin input blots, in triplicate. (D) Quantification of Gα_o_ band intensity, normalized to vinculin band intensity, determined using ImageJ software. Data are means ± SD for three independent experiments. Statistical significance was determined by one-way ANOVA with Dunnett’s multiple comparisons test (α=0.01).

**Figure S2.**
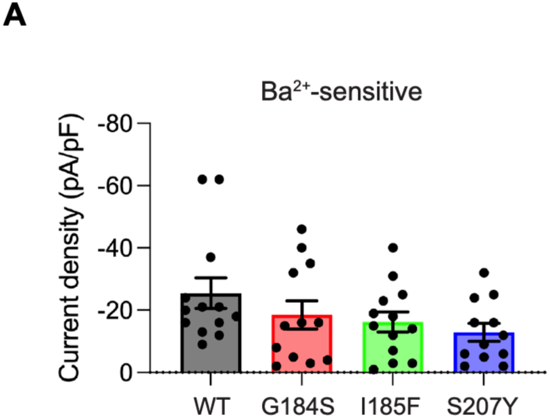
Direct inhibition of GIRK channels by Ba^2+^, as a measure of GIRK channel expression and function. (A) Barium (Ba^2+^)-sensitive GIRK channel currents for Gα_o_ wild-type and representative mutants in HEK293T cells. Data are reported as means ± SEM with 13 independent experiments for wild-type and I185F, 12 independent experiments for G184S and S207Y. Statistical significance was determined by ordinary one-way ANOVA with Dunnett’s multiple comparisons test (α=0.05).

**Figure S3.**
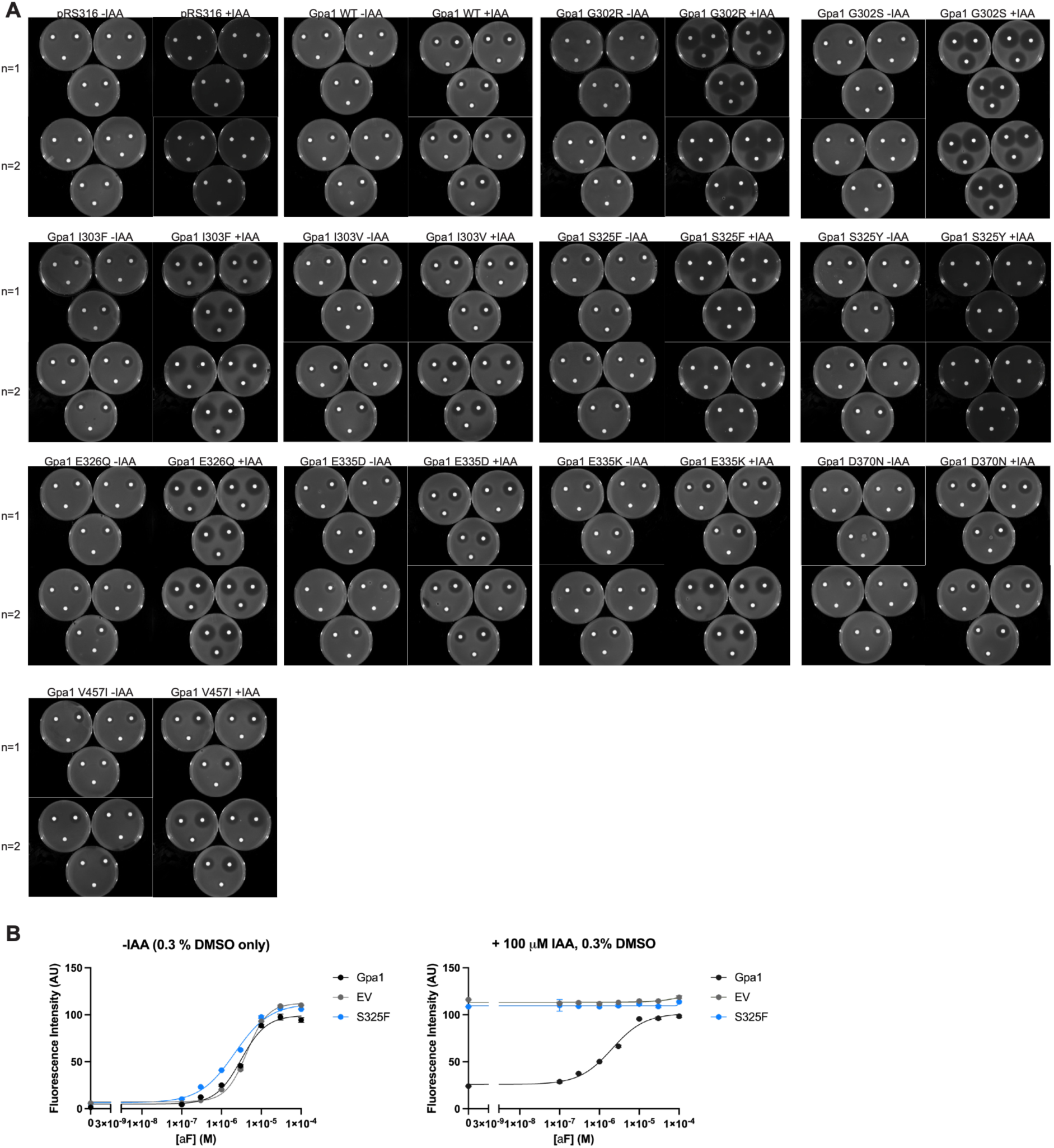
Growth inhibition (halo) assays and unnormalized transcriptional response for Gpa1 wild-type and mutants. (A) Gβγ-induced growth arrest (halo) assays for the tested Gpa1 mutants in yeast. Data are two independent experiments with three measurements each, with or without 10 µM auxin (IAA). (B) Representative unnormalized transcriptional response for wild-type (black), empty vector (gray), and S325F (blue) in the yeast transcription reporter (β-galactosidase) assay. Left; without auxin (DMSO control). Right; with 100 µM auxin. Data are reported as means ± SD and are representative of two independent experiments, three measurements each.

**Figure S4.**
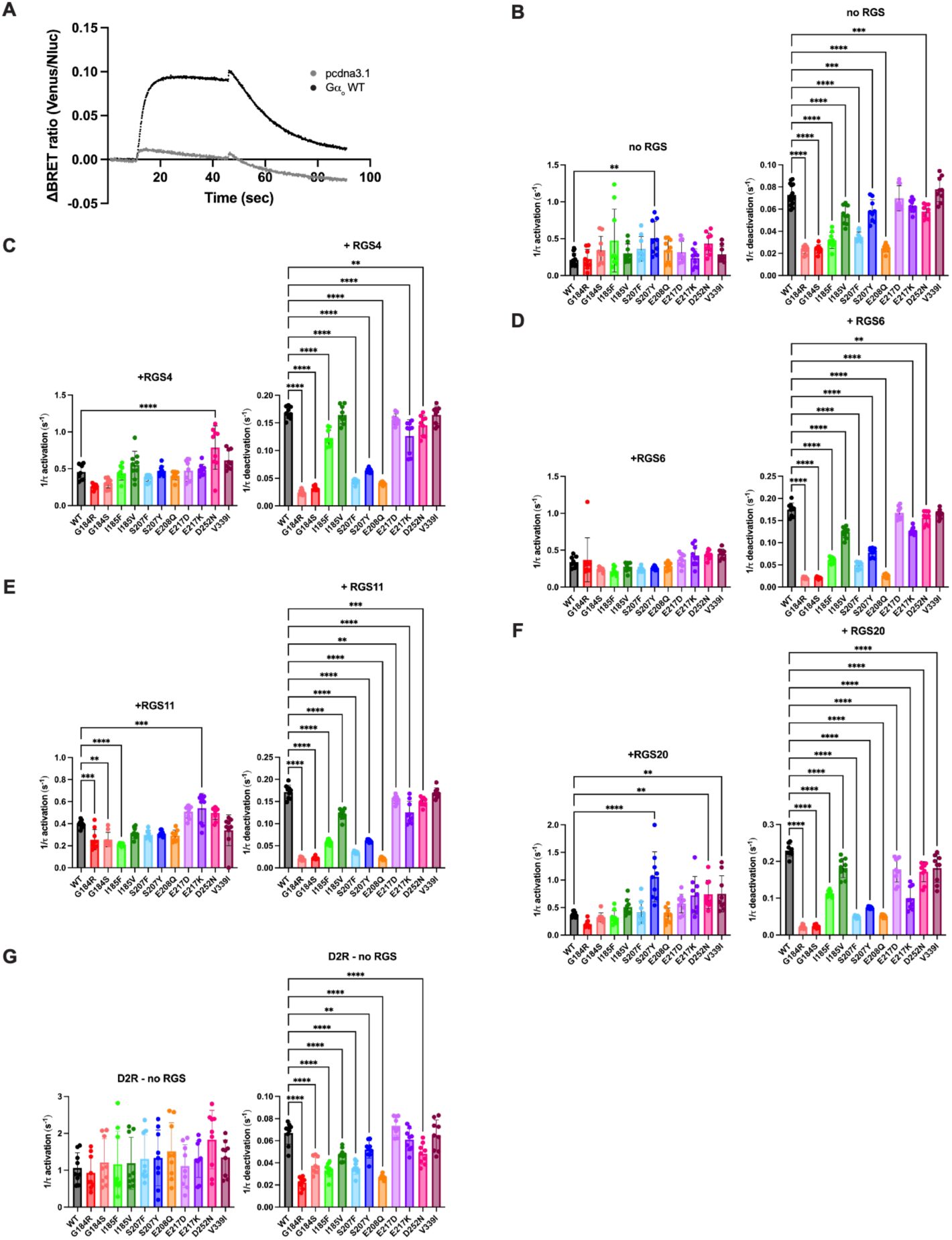
Gβγ-GRK BRET controls and one-way ANOVA results. (A) Baseline-corrected BRET response traces for Gα_o_ wild-type (black) and empty vector (gray) upon activation of the μ-opioid receptor with DAMGO and deactivation with naloxone in HEK293T/17 cells. (B) 1/τ activation rate constants (left) upon μ-opioid receptor stimulation with DAMGO and 1/τ deactivation rate constants (right) after μ-opioid receptor inactivation with naloxone for Gα_o_ wild-type and mutants with no RGS, (C) +RGS4, (D) +RGS6/Gβ_5_, (E) +RGS11/Gβ_5_, and (F) +RGS20. Data are means ± SD from three independent experiments, three measurements each. (G) 1/τ activation rate constants (left) upon dopamine 2 receptor stimulation with dopamine and 1/τ deactivation rate constants (right) after dopamine 2 receptor inactivation with haloperidol for Gα_o_ wild-type and mutants with no RGS. Statistical significance was determined by one-way ANOVA with Dunnett’s multiple comparisons test (α=0.01, **p< 0.01, ***p<0.001, ****p<0.0001). Data are means ± SD from three independent experiments, three measurements each.

**Table S1.** BODIPY-GTP hydrolysis t_1/2_ values for the tested Gα_o_ mutants. Data for Gα_o_ wild-type is reported as mean ± SD for five independent experiments, three measurements each. Data for the Gα_o_ mutants are from two independent experiments, three measurements each. Mutant replicates are reported separately as mean ± SD for one independent experiment, three measurements each. ^1^Previously reported data from (*10*).

| Protein | Bio rep 1<br>$t_{1/2}$ (sec) | Bio rep 2<br>$t_{1/2}$ (sec) | Mean $t_{1/2}$ |
| --- | --- | --- | --- |
| WT | - | - | 58.8 ± 15.3 (n=5) |
| G184R<sup>1</sup> | 97.9 ± 19.0 | 90.6 ± 3.57 | 94.3 |
| G184S | 117 ± 8.04 | 92.5 ± 5.17 | 105 |
| I185F | 59.3 ± 2.32 | 46.9 ± 0.98 | 54.4 |
| I185V<sup>1</sup> | 71.3 ± 10.4 | 79.3 ± 17.6 | 75.3 |
| S207F | 59.6 ± 1.33 | 33.2 ± 1.03 | 46.4 |
| S207Y<sup>1</sup> | 38.1 ± 7.84 | 47.3 ± 0.49 | 42.7 |
| E208Q | 112 ± 6.05 | 66.6 ± 0.44 | 89.4 |
| E217D | 68.2 ± 2.97 | 40.5 ± 2.18 | 54.4 |
| E217K<sup>1</sup> | 58.4 ± 3.36 | 58.4 ± 3.36 | 58.4 |
| D252N<sup>1</sup> | 63.7 ± 2.19 | 60.1 ± 2.94 | 61.9 |
| V339I<sup>1</sup> | 54.6 ± 5.66 | 78.3 ± 17.2 | 66.5 |

**Table S2.** Thermostability melting temperatures (*T_m_*) for the tested Gα_o_ mutants for both nucleotide-bound states. *T_m_* values quantified by fitting a two-state model of thermal unfolding. For Gα_o_ wild-type, *T_m_* values are reported as mean ± SD from five independent experiments, three measurements each. Data for the Gα_o_ mutants are from two independent experiments, three measurements each. Mutant replicates are reported separately as mean ± SD for one independent experiment, three measurements each.

| Protein | Biological replicate 1 |  | Biological replicate 2 |  | Mean |  |
| --- | --- | --- | --- | --- | --- | --- |
| | GDP $T_m$<br>(°C) | GTPγS $T_m$<br>(°C) | GDP $T_m$<br>(°C) | GTPγS $T_m$<br>(°C) | GDP $T_m$<br>(°C) | GTPγS $T_m$<br>(°C) |
| WT | - | - | - | - | 51.3 ± 1.17<br>(n=5) | 57.4 ± 2.38<br>(n=5) |
| G184R<sup>1</sup> | 50.4 ± 0.42 | 56.6 ± 1.04 | 49.7 ± 0.39 | 53.5 ± 1.15 | 50.1 | 55.0 |
| G184S | 49.6 ± 0.41 | 53.8 ± 0.10 | 51.3 ± 0.73 | 55.6 ± 0.47 | 50.5 | 54.7 |
| I185F | 49.7 ± 0.17 | 53.2 ± 0.69 | 51.5 ± 0.58 | 60.0 ± 0.10 | 50.6 | 56.6 |
| I185V<sup>1</sup> | 51.5 ± 0.28 | 57.7 ± 1.24 | 50.8 ± 0.36 | 55.0 ± 0.63 | 51.2 | 56.3 |
| S207F | 51.0 ± 0.17 | 54.2 ± 0.10 | 51.5 ± 0.25 | 55.3 ± 0.35 | 51.2 | 54.8 |
| S207Y<sup>1</sup> | 50.1 ± 2.34 | 56.2 ± 0.36 | 51.2 ± 0.33 | 52.0 ± 0.87 | 50.7 | 54.1 |
| E208Q | 52.1 ± 0.11 | 58.9 ± 0.68 | 51.9 ± 0.55 | 57.0 ± 0.57 | 52.0 | 57.9 |
| E217D | 51.1 ± 0.16 | 58.0 ± 0.40 | 51.4 ± 0.26 | 56.9 ± 0.53 | 51.3 | 57.4 |
| E217K<sup>1</sup> | 52.0 ± 0.33 | 61.6 ± 2.05 | 52.1 ± 0.28 | 55.4 ± 1.17 | 52.1 | 58.5 |
| D252N<sup>1</sup> | 48.0 ± 0.25 | 60.7 ± 0.63 | 48.7 ± 0.40 | 54.1 ± 1.30 | 48.3 | 57.4 |
| V339I<sup>1</sup> | 51.8 ± 1.17 | 56.5 ± 0.49 | 51.0 ± 0.80 | 55.1 ± 0.69 | 51.4 | 55.8 |

**Table S3.** EC_50_ values for all tested Gpa1 mutants treated with auxin. For Gpa1 wild-type, EC_50_s are reported as mean ± SD from fifteen independent experiments, three measurements each. Data for the Gpa1 mutants are from two independent experiments, three measurements each. Mutant replicates are reported separately as mean ± SD for one independent experiment, three measurements each. “N.M.”, not measurable.

| Protein | Bio rep 1<br>EC<sub>50</sub> (μM) | Bio rep 2<br>EC<sub>50</sub> (μM) | Mean EC<sub>50</sub> (μM) |
| --- | --- | --- | --- |
| WT | - | - | 2.82 ± 0.54 (n=15) |
| G302R | 0.07 ± 0.02 | 0.07 ± 0.01 | 0.07 |
| G302S | 0.13 ± 0.03 | 0.18 ± 0.02 | 0.15 |
| I303F | 1.29 ± 0.03 | 1.08 ± 0.11 | 1.19 |
| I303V | 2.30 ± 0.05 | 2.11 ± 0.13 | 2.21 |
| S325F | N.M. | N.M. | N.M. |
| S325Y | 3.37 ± 0.21 | 3.49 ± 0.14 | 3.43 |
| E364Q | 0.30 ± 0.05 | 0.30 ± 0.06 | 0.30 |
| E335D | 2.15 ± 0.09 | 2.72 ± 0.21 | 2.43 |
| E335K | 1.67 ± 0.11 | 1.86 ± 0.16 | 1.76 |
| D370N | 4.40 ± 0.42 | 4.12 ± 0.10 | 4.26 |
| V457I | 3.18 ± 0.19 | 4.42 ± 0.23 | 3.80 |

**Table S4.** *k*_app_ values for Gα_o_ wild-type and mutants. *k*_app_ values are calculated by fitting a one-phase decay curve fit to the deactivation traces from the kinetic BRET experiments. Data are means ± SD of three independent experiments, three measurements each.

| Protein | -RGS ($s^{-1}$) | +RGS4 ($s^{-1}$) | +RGS6/$G\beta_5$ ($s^{-1}$) | +RGS11/$G\beta_5$ ($s^{-1}$) | +RGS20 ($s^{-1}$) |
| --- | --- | --- | --- | --- | --- |
| WT | $0.073 \pm 0.009$ | $0.169 \pm 0.011$ | $0.175 \pm 0.015$ | $0.172 \pm 0.015$ | $0.230 \pm 0.017$ |
| G184R | $0.023 \pm 0.004$ | $0.024 \pm 0.005$ | $0.021 \pm 0.002$ | $0.020 \pm 0.004$ | $0.022 \pm 0.005$ |
| G184S | $0.024 \pm 0.004$ | $0.032 \pm 0.004$ | $0.021 \pm 0.002$ | $0.022 \pm 0.003$ | $0.023 \pm 0.004$ |
| I185F | $0.032 \pm 0.007$ | $0.123 \pm 0.015$ | $0.061 \pm 0.007$ | $0.059 \pm 0.005$ | $0.113 \pm 0.008$ |
| I185V | $0.055 \pm 0.008$ | $0.165 \pm 0.016$ | $0.124 \pm 0.012$ | $0.121 \pm 0.012$ | $0.182 \pm 0.026$ |
| S207F | $0.035 \pm 0.005$ | $0.044 \pm 0.004$ | $0.050 \pm 0.008$ | $0.035 \pm 0.003$ | $0.050 \pm 0.003$ |
| S207Y | $0.058 \pm 0.011$ | $0.064 \pm 0.004$ | $0.078 \pm 0.010$ | $0.061 \pm 0.003$ | $0.074 \pm 0.003$ |
| E208Q | $0.025 \pm 0.004$ | $0.041 \pm 0.003$ | $0.026 \pm 0.004$ | $0.021 \pm 0.003$ | $0.051 \pm 0.004$ |
| E217D | $0.070 \pm 0.011$ | $0.157 \pm 0.001$ | $0.167 \pm 0.014$ | $0.154 \pm 0.009$ | $0.178 \pm 0.034$ |
| E217K | $0.063 \pm 0.006$ | $0.126 \pm 0.029$ | $0.127 \pm 0.008$ | $0.125 \pm 0.025$ | $0.010 \pm 0.031$ |
| D252N | $0.058 \pm 0.006$ | $0.146 \pm 0.018$ | $0.156 \pm 0.015$ | $0.149 \pm 0.010$ | $0.172 \pm 0.027$ |
| V339I | $0.078 \pm 0.011$ | $0.165 \pm 0.016$ | $0.165 \pm 0.010$ | $0.169 \pm 0.011$ | $0.182 \pm 0.038$ |

**Table S5.** k_GAP_ values for Gα_o_ wild-type and mutants. *k*_GAP_ values are calculated by subtracting k_app_ (no RGS) from k_app_ (+RGS) for a given mutant. Data are means ± SD of three independent experiments, three measurements each.

| Protein | +RGS4 ($\text{s}^{-1}$) | +RGS6/$\text{G}\beta_5$ ($\text{s}^{-1}$) | +RGS11/$\text{G}\beta_5$ ($\text{s}^{-1}$) | +RGS20 ($\text{s}^{-1}$) |
| --- | --- | --- | --- | --- |
| WT | $0.096 \pm 0.014$ | $0.102 \pm 0.018$ | $0.099 \pm 0.018$ | $0.157 \pm 0.019$ |
| G184R | $0.001 \pm 0.006$ | $-0.003 \pm 0.004$ | $-0.003 \pm 0.005$ | $-0.002 \pm 0.006$ |
| G184S | $0.008 \pm 0.005$ | $-0.003 \pm 0.004$ | $-0.002 \pm 0.005$ | $-0.001 \pm 0.006$ |
| I185F | $0.091 \pm 0.016$ | $0.029 \pm 0.010$ | $0.027 \pm 0.009$ | $0.081 \pm 0.011$ |
| I185V | $0.110 \pm 0.018$ | $0.069 \pm 0.015$ | $0.066 \pm 0.014$ | $0.127 \pm 0.027$ |
| S207F | $0.009 \pm 0.006$ | $0.015 \pm 0.009$ | $0.000 \pm 0.005$ | $0.015 \pm 0.006$ |
| S207Y | $0.006 \pm 0.012$ | $0.020 \pm 0.014$ | $0.002 \pm 0.011$ | $0.016 \pm 0.011$ |
| E208Q | $0.016 \pm 0.004$ | $0.001 \pm 0.005$ | $-0.004 \pm 0.004$ | $0.026 \pm 0.005$ |
| E217D | $0.087 \pm 0.015$ | $0.098 \pm 0.018$ | $0.085 \pm 0.014$ | $0.108 \pm 0.036$ |
| E217K | $0.064 \pm 0.030$ | $0.064 \pm 0.010$ | $0.062 \pm 0.026$ | $0.037 \pm 0.031$ |
| D252N | $0.088 \pm 0.019$ | $0.099 \pm 0.016$ | $0.091 \pm 0.012$ | $0.114 \pm 0.028$ |
| V339I | $0.087 \pm 0.020$ | $0.088 \pm 0.015$ | $0.092 \pm 0.016$ | $0.104 \pm 0.040$ |

**Table S6.**
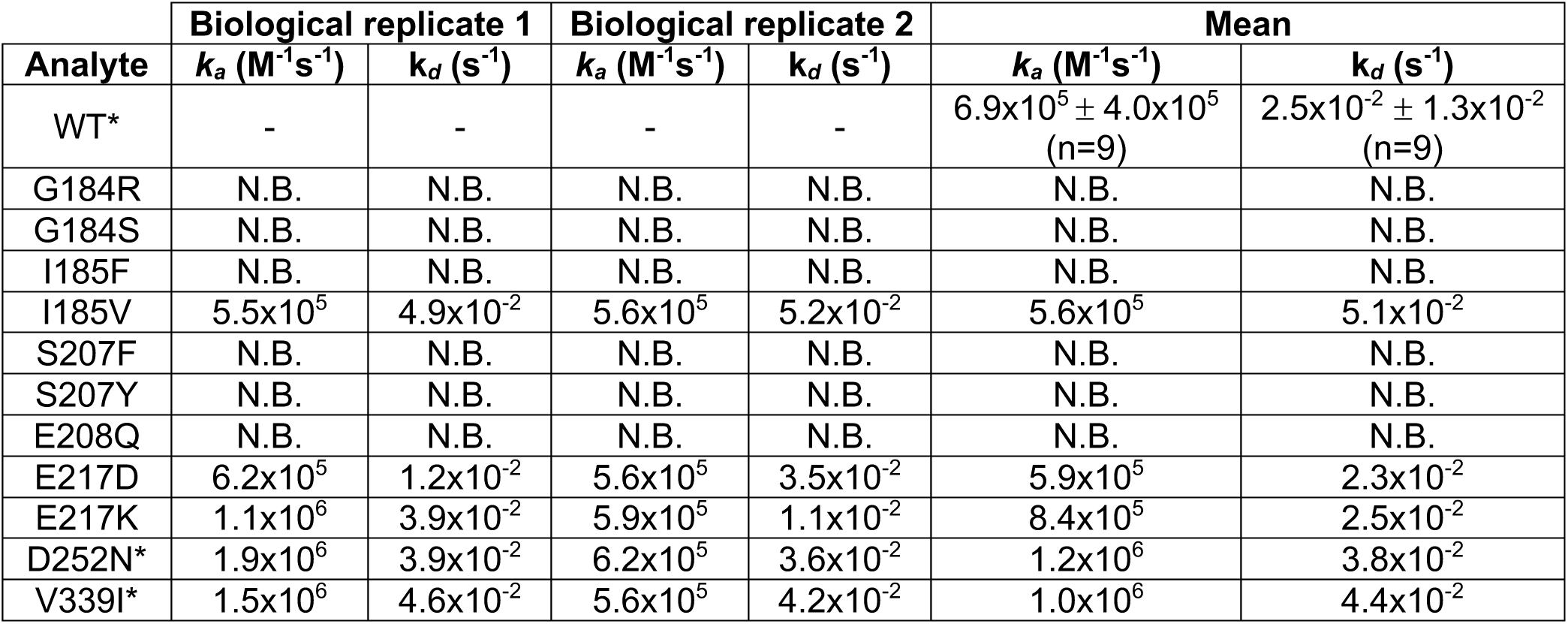
Kinetic values for SPR experiments. Data are means ± SD of 9 biological replicates for Gα_o_ wild-type, 2 biological replicates for Gα_o_ mutants. The *K*_D_, *k_a_*, and *k_d_* values were generated by the Biacore 8K+ Insight Evaluation Software. “N.B.”, no binding. *For two WT, one D252N, and one V339I replicates, values were derived by removing reference subtraction to correct for poor fits due to mismatches between the reference and active channels.

| Analyte | Biological replicate 1 |  | Biological replicate 2 |  | Mean |  |
| --- | --- | --- | --- | --- | --- | --- |
| | $k_a$ ($M^{-1}s^{-1}$) | $k_d$ ($s^{-1}$) | $k_a$ ($M^{-1}s^{-1}$) | $k_d$ ($s^{-1}$) | $k_a$ ($M^{-1}s^{-1}$) | $k_d$ ($s^{-1}$) |
| WT* | - | - | - | - | $6.9 \times 10^5 \pm 4.0 \times 10^5$<br>(n=9) | $2.5 \times 10^{-2} \pm 1.3 \times 10^{-2}$<br>(n=9) |
| G184R | N.B. | N.B. | N.B. | N.B. | N.B. | N.B. |
| G184S | N.B. | N.B. | N.B. | N.B. | N.B. | N.B. |
| I185F | N.B. | N.B. | N.B. | N.B. | N.B. | N.B. |
| I185V | $5.5 \times 10^5$ | $4.9 \times 10^{-2}$ | $5.6 \times 10^5$ | $5.2 \times 10^{-2}$ | $5.6 \times 10^5$ | $5.1 \times 10^{-2}$ |
| S207F | N.B. | N.B. | N.B. | N.B. | N.B. | N.B. |
| S207Y | N.B. | N.B. | N.B. | N.B. | N.B. | N.B. |
| E208Q | N.B. | N.B. | N.B. | N.B. | N.B. | N.B. |
| E217D | $6.2 \times 10^5$ | $1.2 \times 10^{-2}$ | $5.6 \times 10^5$ | $3.5 \times 10^{-2}$ | $5.9 \times 10^5$ | $2.3 \times 10^{-2}$ |
| E217K | $1.1 \times 10^6$ | $3.9 \times 10^{-2}$ | $5.9 \times 10^5$ | $1.1 \times 10^{-2}$ | $8.4 \times 10^5$ | $2.5 \times 10^{-2}$ |
| D252N* | $1.9 \times 10^6$ | $3.9 \times 10^{-2}$ | $6.2 \times 10^5$ | $3.6 \times 10^{-2}$ | $1.2 \times 10^6$ | $3.8 \times 10^{-2}$ |
| V339I* | $1.5 \times 10^6$ | $4.6 \times 10^{-2}$ | $5.6 \times 10^5$ | $4.2 \times 10^{-2}$ | $1.0 \times 10^6$ | $4.4 \times 10^{-2}$ |

**Table S7.**
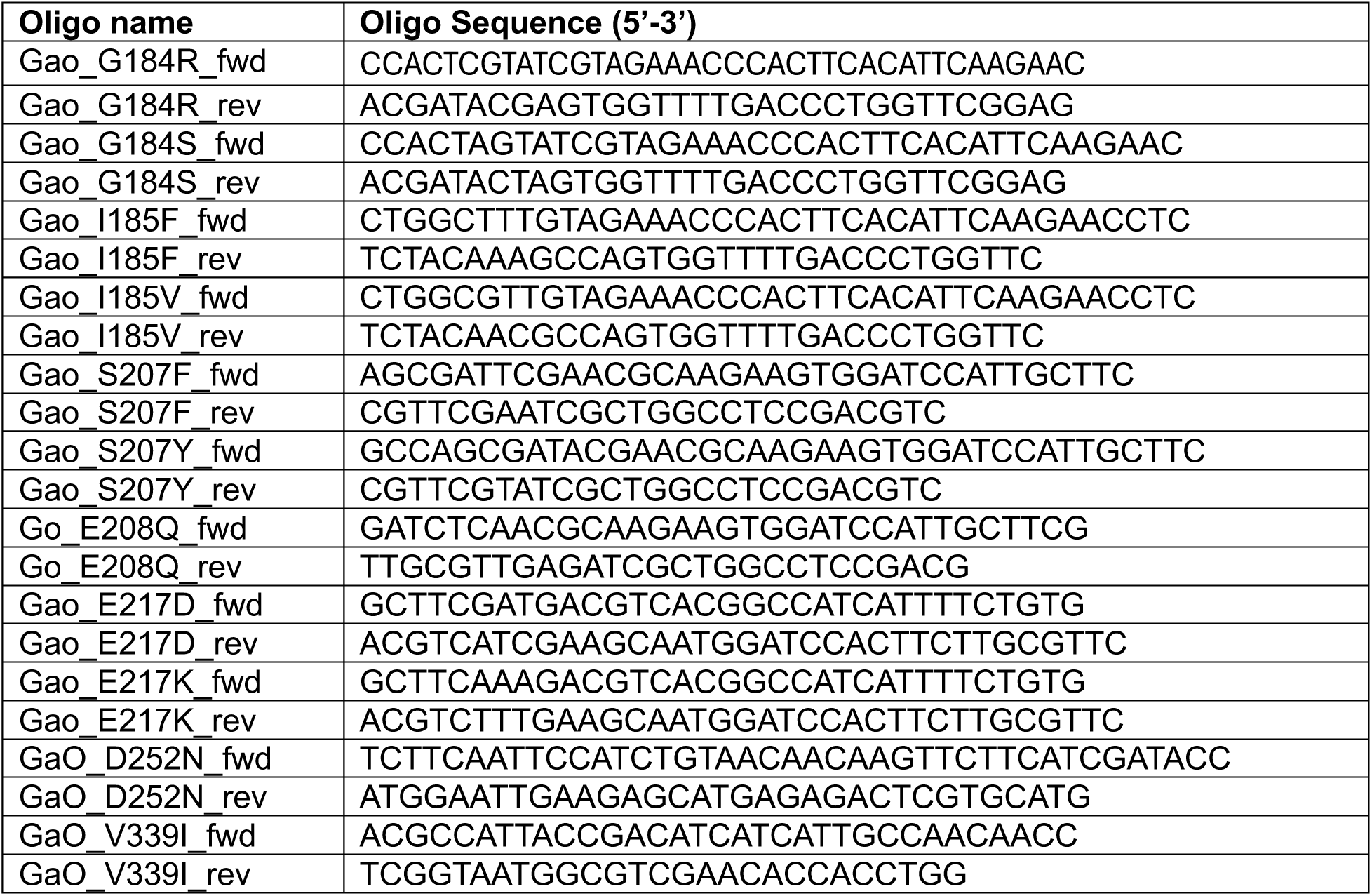
Mutagenesis oligonucleotide sequences.

